# Cognitive state dependent enhancement of cognitive control with transcranial magnetic stimulation

**DOI:** 10.64898/2025.12.12.694061

**Authors:** Aaron N. McInnes, Victoria L. Pipia, Kiyana L. Maynard, Güldamla Kalender, Alik S. Widge

## Abstract

**Background:** Repetitive transcranial magnetic stimulation (rTMS) is an effective treatment for major depressive disorder (MDD), yet variability of therapeutic responses remains high. A key contributor to this variability may be state-dependent effects of brain stimulation, where activity in underlying circuits can shape the propagation of TMS-evoked activity. Thus, constraining the target circuits’ state by engaging cognition may make TMS’ effects more consistent.

**Objective:** We tested whether TMS’ effects on cognitive control, a transdiagnostic construct implicated across psychiatric disorders and a putative mediator of TMS efficacy, are state-dependent.

**Methods:** Participants (N = 25) completed a context-dependent behavioural assay of cognitive control before and after we delivered rTMS to the prefrontal cortex (PFC). During rTMS, participants performed either the cognitive control task (active-state TMS), or performed a context-independent perceptual task (control-state TMS). We assessed changes in downstream behavioural metrics of circuit function, as well as neural indices of cognitive control measured from electroencephalography (EEG).

**Results:** rTMS enhanced downstream readouts of cognitive control only when stimulation was delivered concurrent with the cognitive control task. Similarly, only active-state TMS modulated theta band oscillatory activity, which is thought to be a marker of cognitive control engagement

**Conclusion:** These findings demonstrate that TMS’ effects on cognitive control are state-dependent, in which endogenous engagement of PFC-anchored networks can shape the magnitude and functional relevance of TMS-induced plasticity. Considering cognitive states during TMS may therefore offer a framework to enhance and/or accelerate TMS’ therapeutic effects.

---

Repetitive transcranial magnetic stimulation (rTMS) is a non-invasive form of brain stimulation with established efficacy for treatment-resistant major depressive disorder (MDD)^1^. TMS is thought to evoke neuroplastic changes in the stimulated region and connected neural circuitry, altering aberrant network activity underlying the symptoms of psychiatric disorders^2,3^. TMS is safe and effective^4^, but the magnitude and durability of clinical responses are highly variable^5–10^. Thus, identifying factors that govern TMS’ effective engagement of its neural targets is a central challenge in improving clinical outcomes.

A candidate source of therapeutic variability is the dependence of neuromodulation on endogenous neural activity at the time of stimulation (i.e., state-dependence)^11–17^. Consistent with Hebbian^18^ and metaplasticity^19^ models of neural learning, circuits that are already active may be in a more biophysically permissive state for modification, making them prime targets for intervention-induced plasticity. In line with this notion, functional reorganization of sensorimotor cortex by neuromodulation depends on the underlying network structure of the brain^20^. Similarly, antidepressant responses to TMS may depend on functional connectivity of underlying networks^21^. However, it remains unclear how specific manipulations of underlying circuit activity can shape the magnitude and functional specificity of TMS for MDD.

Consideration of state-dependence has motivated efforts to combine TMS with psychotherapy^22–24^, and one approved indication depends on this approach^25^. However, logistical barriers^26^ and variability in patients’ engagement with psychotherapy exercises may limit this approach^24^. An alternative strategy is to manipulate the underlying circuit state through computerised cognitive exercises. The propagation of TMS-evoked activity depends on underlying function in sensorimotor^16,27–29^, attention^30–33^, and memory^34–36^ networks. This input selectivity raises the possibility that actively engaging target neural circuits may sensitise them to neuromodulation, while sparing quiescent non-target circuits^12^.

Cognitive control provides a principled framework to engage the circuits commonly targeted in TMS interventions for depression. Control involves the suppression of prepotent responses in favour of more adaptive ones, and depends on distributed brain networks anchored to the prefrontal cortex (PFC)^37^. PFC, in turn, is the most common target of TMS for MDD, and TMS may in part work by inducing PFC neuroplasticity that supports cognitive control^38,39^.

Conversely, if a TMS recipient is exerting cognitive control while stimulation is delivered, that endogenous activity might interact with and amplify the plasticity effect. Thus, by manipulating the engagement of cognitive control mechanisms, it becomes possible to test how the activation of those PFC-anchored networks shapes the magnitude, topology, and functional relevance of TMS-evoked activity.

Here, we tested whether the cognitive-behavioural and neural effects of rTMS are state-dependent. rTMS was applied to the PFC while participants performed either a context-dependent cognitive control task, or a context-independent perceptual discrimination task. rTMS modified cognitive control performance and neural indices of control only when the underlying circuits were active during stimulation. Considering and controlling cognitive states may therefore offer a framework to enhance and/or accelerate TMS’ therapeutic effects.

## 2. Materials and Methods

### 2.1. Participants

Twenty-five individuals free from potential contraindications to TMS^40^ participated (14 female, mean age = 27.65 years, SD = 9.67, range = 19 – 65). We also excluded individuals who self-reported any psychiatric diagnoses so that we could characterise typical state-dependent brain responses without influence from aberrant dorsolateral PFC (and related network) activity. For our planned primary analysis of behavioural modification by rTMS as a function of the behavioural state at the time of stimulation, our sample size was determined via power analysis of (generalised) linear mixed effects models^41,42^ to achieve 80% statistical power and targeting a moderate effect size (η_p_ ^2^ = 0.08) based on DLPFC-neuromodulation studies in cognition and executive function^43,44^. The protocol was approved by the local institutional review board and all participants provided informed, written consent in accordance with the declaration of Helsinki.

### 2.2. Procedures

To examine state dependence of prefrontal rTMS, we applied trains of rTMS at 100% of resting motor threshold during either a PFC-dependent cognitive control task, or a perceptual discrimination task that should be less PFC-dependent, across two separate sessions, approximately one week apart, counterbalanced across participants. A detailed description of the stimulation parameters is provided in the **Supplementary Materials**. For the duration of the experiment, we recorded EEG at 1 kHz (Brain Products actiCHamp, GmbH, Munich, Germany) with 64 active electrodes in the international 10-20 arrangement^45^ and with a ground electrode placed at the FPz site.

#### 2.2.1. Behavioural tasks

##### 2.2.1.1. Translational Orientation Pattern Expectancy

To measure cognitive control function (Pre-rTMS and Post-rTMS), and engage cognitive control circuitry (ActiveState_rTMS_), participants completed the translational orientation pattern expectancy (TOPX) task^46^. The task presents two visual stimuli, a cue and a probe, each requiring a behavioural response. A frequent cue-probe target sequence produces a prepotent response tendency that must be countermanded in infrequent non-target sequences (**Figure 1**).

**Figure 1.**
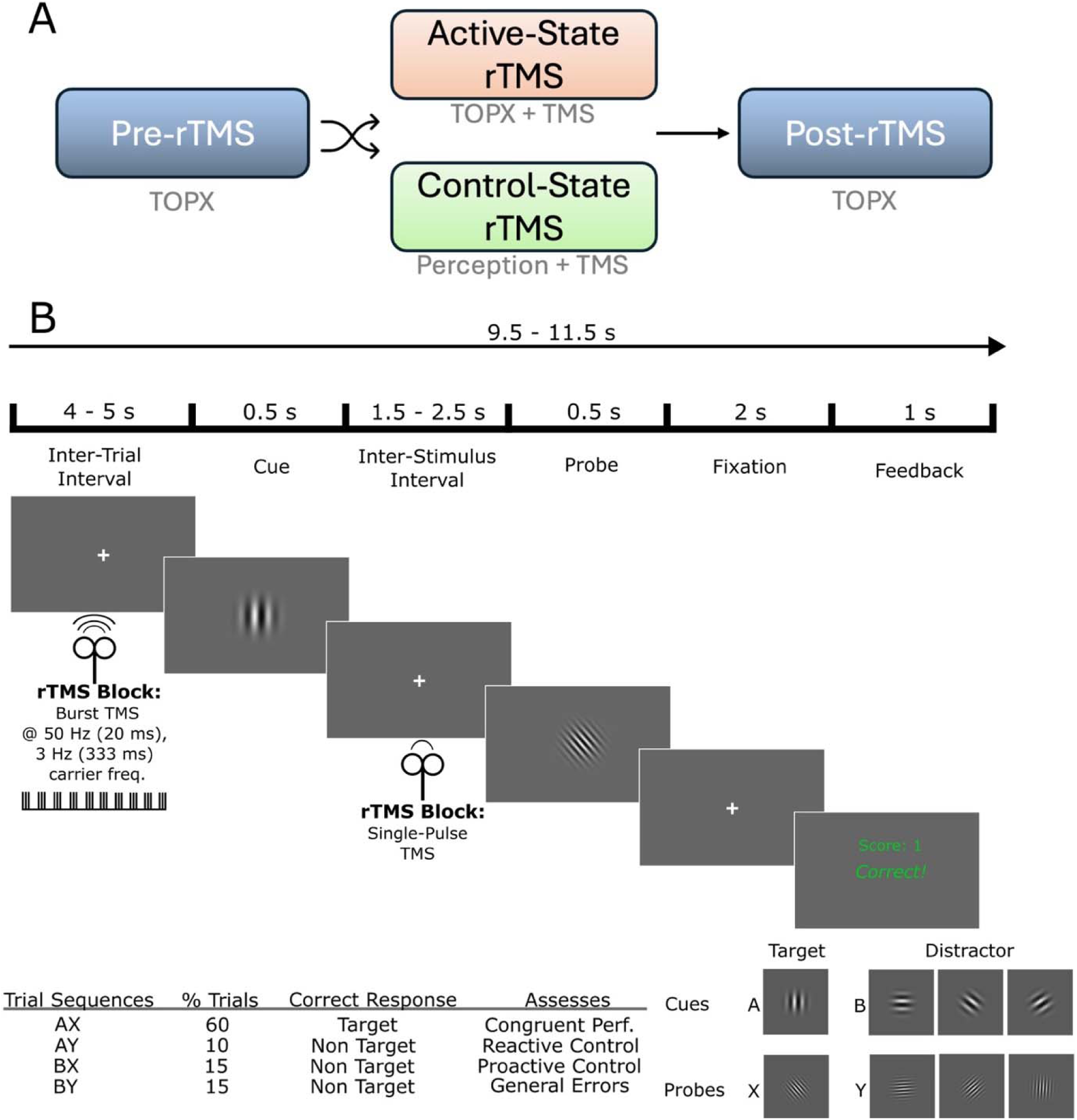
Sequence of events during the translational orientation pattern expectancy (TOPX) and perceptual tasks. (A) Block order of the experiment, in which participants complete a Pre- and Post-rTMS assessment of cognitive control, without TMS administered. In the rTMS block, the behavioural state is manipulated. In active-state rTMS, participants complete the context-dependent TOPX task while receiving rTMS. In the control-state rTMS condition, participants complete a perceptual task which is context-independent while receiving rTMS. Participants complete both conditions, order counterbalanced, with a one-week washout period. (B) Sequence of events in the TOPX task. A cue and probe stimulus appear. Participants must give one response when a target (AX) sequence is presented, and an alternate response when a non-target sequence (AY, BX, BY) sequence is presented. In the perceptual task, the same stimuli are presented, but the contextual goal changes; participants respond to only the orientation of each Gabor stimulus. In the rTMS block, trains of rTMS were delivered during the inter-trial interval of TOPX (active-state TMS) or Perceptual (control-state TMS) tasks. We also administered single-pulse TMS during the cue-probe inter-stimulus interval of the TOPX and Perceptual tasks to assess cortical excitation and reactivity to TMS.

Participants completed TOPX before (baseline measure) and after (change measure) receiving rTMS. Every run of the task consisted of four blocks of 30 trials. Participants completed 16 practice trials prior to beginning experimental trials of TOPX. In the ActiveState_rTMS_ condition, participants also completed a run of TOPX while receiving trains of prefrontal rTMS.

##### 2.2.1.2. Perceptual Task

In the ControlState_rTMS_ condition, participants completed a perceptual discrimination task while receiving rTMS. The perceptual task had the same stimulus parameters and trial structure as TOPX, but did not require context processing or inhibition of prepotent responses. Instead, participants responded according to the orientation of each stimulus. See **Supplementary Materials** for a description of the task parameters in the TOPX and perceptual tasks.

### 2.3. Data Processing and Statistical Analyses

#### 2.3.1. Behavioural Data

Statistical analysis was performed in R^47^. Two participants did not complete the second experimental session and were excluded from analysis. For each TOPX run, we computed accuracy, probe reaction time (RT), D′-Context, and proactive behavioural control (PBI) indices, and quantified pre–post rTMS change using metric-appropriate difference or ratio scores (definitions in **Supplementary Materials**). Those change scores were analysed as a function of behavioural state at the time of TMS (ActiveState_rTMS_, ControlState_rTMS_) using a series of linear mixed-effects models, including session number (1, 2) as a fixed effect and subject IDs as a random intercept. Given their positive, right-skewed distribution, we analysed RTs using a generalised linear mixed effects model with a gamma link function and identity distribution, as in our past work^48–50^.

Recent work from our group^49,51–53^ and others^54–57^ suggests that neurostimulation of PFC and connected structures affects evidence accumulation processes. To assess those after our task-linked intervention, we hierarchically fit drift diffusion models (DDMs) to reaction times and response accuracy to infer latent processes of evidence accumulation towards decisions in TOPX. Model specification followed prior work applying hierarchical DDMs to expectancy-based tasks^57^, where the *v* and *a* parameters were permitted to vary by trial sequence (AX, AY, BX, BY), while *z* was permitted to vary only by cue type (A Cue, B Cue), and *t* was held constant across conditions and estimated hierarchically with a group-level prior. We verified that this model structure fit the data well using an information criterion analysis (Supplementary Methods).

#### 2.3.2. Electroencephalography

EEG data were pre-processed offline using EEGLAB^58^ and FieldTrip^59^ toolboxes in MATLAB (R2023a, Mathworks, USA). EEG data from -2 to +15 ms around TMS pulses were removed and interpolated to mitigate high amplitude TMS pulse artifacts, as recommended in standard TMS-EEG procedures^60^. Data were resampled to 500 Hz and any channels showing poor signal were identified based on kurtosis and interpolated. We applied a two-step independent component analysis (ICA) procedure to first remove TMS-evoked pulse and muscle artifacts, and a second ICA to remove remaining artifactual components^60,61^. In addition, we band-pass filtered at 1 – 50 Hz and re-referenced to the average of all channels.

To extract task-evoked and TMS-evoked oscillatory activity, we applied a Morlet wavelet decomposition on single trials^62^. Wavelets had frequency-dependent widths from 3 – 50 Hz, starting at 3 cycles at 3 Hz and adding 0.8 cycles for each 1 Hz. Event-related spectral perturbations (ERSPs) were computed by applying a pre-stimulus divisive baseline in the linear power domain from -500 to -100 ms relative to the target event.

We also considered that cognitive control is not a unitary construct. The Dual Mechanisms of Control (DMC) framework describes two distinct modes of proactive versus reactive control, each supported by partially dissociable PFC-based networks^63^. The TOPX task, which is based upon the AX-Continuous Performance Test (AX-CPT^64–67)^ probes both these mechanisms by modifying the type of contextual information provided prior to a context-dependent response^65,68^. For instance, distractor cues provide sufficient contextual information to indicate that the trial sequence is a non-target, allowing participants to maintain goal-relevant information in advance and deploy proactive control. In contrast, target cues provide insufficient context to determine the appropriate response until presentation of the probe stimulus, thereby evoking greater reactive control demands (**Figure 1**). Because proactive and reactive trials were expected to engage partially distinct spatial distributions of activity^69^, we analysed ERSPs using cluster-based permutation tests to identify scalp-wide, data-driven regions of interest, rather than imposing a-priori electrode or source locations. For task-evoked ERSPs, we analysed the difference between blocks (post-rTMS – pre-rTMS) as a function of behavioural states during rTMS (cognitive control, perception), and for each stimulus type (target, distractor). Theta band activity has a known role in executive functioning^70–73^, and thus, we primarily expected TMS to modulate activity within that band during the cognitive control task. Therefore, we tested for electrode clusters within the theta band (4 – 8 Hz) and within the time window 0 – 1000 ms relative to the task stimulus, applying a threshold for cluster formation of at least two electrodes. To test for spectral specificity (as opposed to broadband effects) within the derived spatially defined effects, we then performed follow-up time-frequency cluster-based permutation tests restricted to the averaged activity across electrodes in each significant cluster. We applied similar tests for TMS-evoked ERSPs, but compared the difference between the cognitive control and perceptual tasks, using a time window of 0 – 500 ms^74^ relative to single-pulse TMS or the end of the rTMS train.

## 3. Results

### 3.1. State-Dependent Enhancement of Cognitive Control

#### 3.1.1. Accuracy

Task-linked rTMS enhanced cognitive control, but only when the target circuits were engaged by the task (F_(1, 45)_ = 6.11, *p* = .017, *R²* = .121), with ratios of accuracy significantly greater than μ = 1 (baseline) for ActiveState_rTMS_ (*M* = 1.03, *SD* = 0.04, *p* = .005), but not for ControlState_rTMS_ (*M* = 1, *SD* = 0.04, *p* = .985; **Figure 2A**). Raw behavioural data are presented in the **Supplementary Materials Figure S1**.

**Figure 2.**
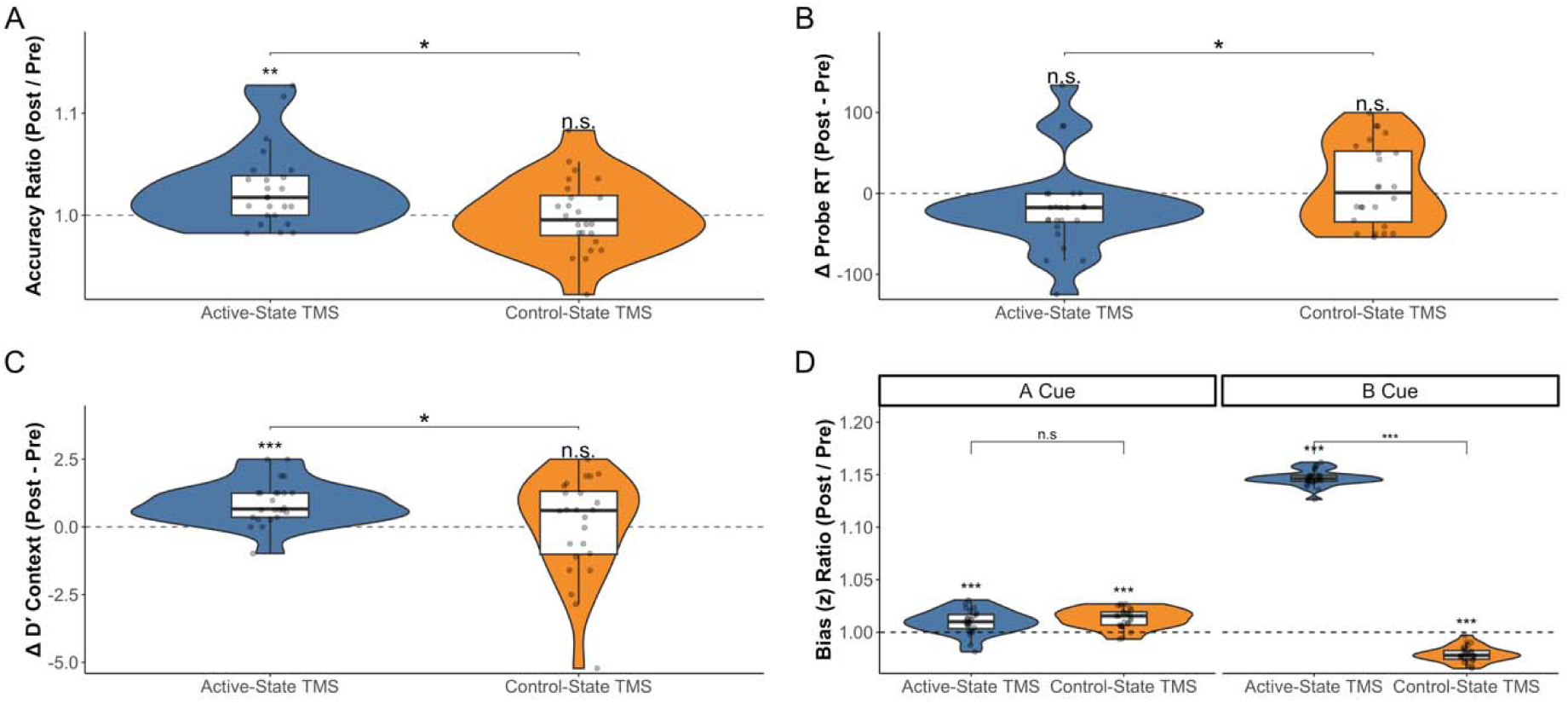
Enhancement of cognitive control by active-state TMS. (A) Accuracy was enhanced by TMS when cognitive control circuitry was engaged, but not when non-target circuits were engaged during TMS. (B) Probe RTs were numerically shortened by Active-State TMS, but that difference was not significantly different from μ = 0 (no change). However, RT shortening showed a significant improvement after Active-State TMS relative to Control-State TMS. (C) Contextual processing, represented by D’-Context, showed a similar state-dependent enhancement after Active-State TMS, but not Control-State TMS. (D) Evidence accumulation bias estimated by drift diffusion modelling showed a state dependent modulation by TMS, particularly for non-target cues, whereby starting point bias was increased in Active-State TMS, but decreased in the Control-State TMS condition. Violins depict probability density distributions for each condition; boxplots depict the between-subject median and interquartile range for each condition; points represent individual-subject change scores. Significance (FDR-corrected) is denoted above each condition with reference to mean difference = 0 or mean ratio = 1 (no change). Significance (FDR-corrected) is denoted between conditions determined by pairwise contrasts. *** = *p* < .001, ** = *p* < .01, * = *p* < .05, n.s = *p* > .05.

#### 3.1.2. Reaction Time

Reaction time was differentially shortened after TMS as a function of rTMS State (F_(1, 22.39)_ = 6.70, *p* = .017, *R²* = .099; **Figure 2B**), and indicated a numeric shortening of reaction time in the ActiveState_rTMS_ condition (*M* = -17.49 ms, *SD* = 55.14), but not the ControlState_rTMS_ condition (*M* = 14.52, *SD* = 51.52). Those RT differences were not significantly different from μ = 0 (no change) in either ActiveState_rTMS_ or ControlState_rTMS_ conditions, but the difference between RTs from the two conditions (ActiveState_rTMS_ relative to ControlState_rTMS_) was significant (*p* = .017).

#### 3.1.3. Contextual Behavioural Performance

Contextual processing as indexed by D -Context showed a state dependent enhancement following TMS (*F_(1, 45)_* = 4.62, *p* = .037, *R²* = .094; **Figure 2C**), with significantly greater TMS-evoked change for ActiveState_rTMS_ (*M* = 0.88, *SD* = 0.80, *p* < .001) relative to μ = 0 (p < .001) and relative to ControlState_rTMS_ (*p* = .043). In contrast, ControlState_rTMS_ differences were not significantly different from μ = 0 (*M* = 0.02, *SD* = 1.83, *p* = .965). PBI measures did not significantly change after TMS (see **Supplementary Materials**).

#### 3.1.4. Evidence accumulation

Hierarchical drift diffusion modelling revealed no modulation of drift rate or decision threshold. In contrast, starting point bias showed a state-dependent modulation by rTMS which depended on contextual information (cue type with rTMS state interaction on bias ratios, *F_(1, 68.48)_* = 2334.45, *p* < .001, *R²* = .962; **Figure 2D**). For A cues, the rTMS evoked change in starting point bias was significantly greater than μ = 1 for both the ActiveState_rTMS_ (*M* = 1.01, *SD* = 0.01, *p* < .001) and ControlState_rTMS_ (*M* = 1.01, *SD* = 0.01, *p* < .001) conditions, but the change was not significantly different between the two conditions (*p* = .259). For B cues, the rTMS evoked change in starting point bias showed a significant increase relative to μ = 1 for ActiveState_rTMS_ (*M* = 1.15, *SD* = 0.01, *p* < .001), a significant decrease relative to μ = 1 for ControlState_rTMS_ (*M* = 0.98, *SD* = 0.01, *p* < .001), and the change was significantly different between the two conditions (*p* < .001). In a follow-up experiment, we found that task-switching or practice effects do not explain our observed behavioural effects (see **Supplementary Materials**).

### 3.2. State Dependent Modulation of Theta Band Activity

#### 3.2.1. Cortical Engagement by TMS Depends on Cognitive State

We examined state dependence of TMS’ cortical engagement by comparing theta band ERSPs evoked after single-pulse and repetitive TMS for each of the cognitive control and perceptual tasks. Cluster-based tests did not identify statistically significant electrode clusters that differed as a function of the behavioural state during stimulation. However, we tested for differences in theta band ERSPs specifically at the stimulation site (F3) and found that single-pulse TMS-evoked ERSPs were on average 0.58 dB larger in the cognitive control task in comparison to the perceptual one and that difference was statistically significant against a null distribution generated by shuffled task labels (*p* = .014). Similarly, following the rTMS trains, theta band ERSPs were 0.49 dB larger on average during the cognitive control task relative to the perceptual one (*p* = .021). ERSPs evoked by TMS are shown in **Figure 3**.

**Figure 3.**
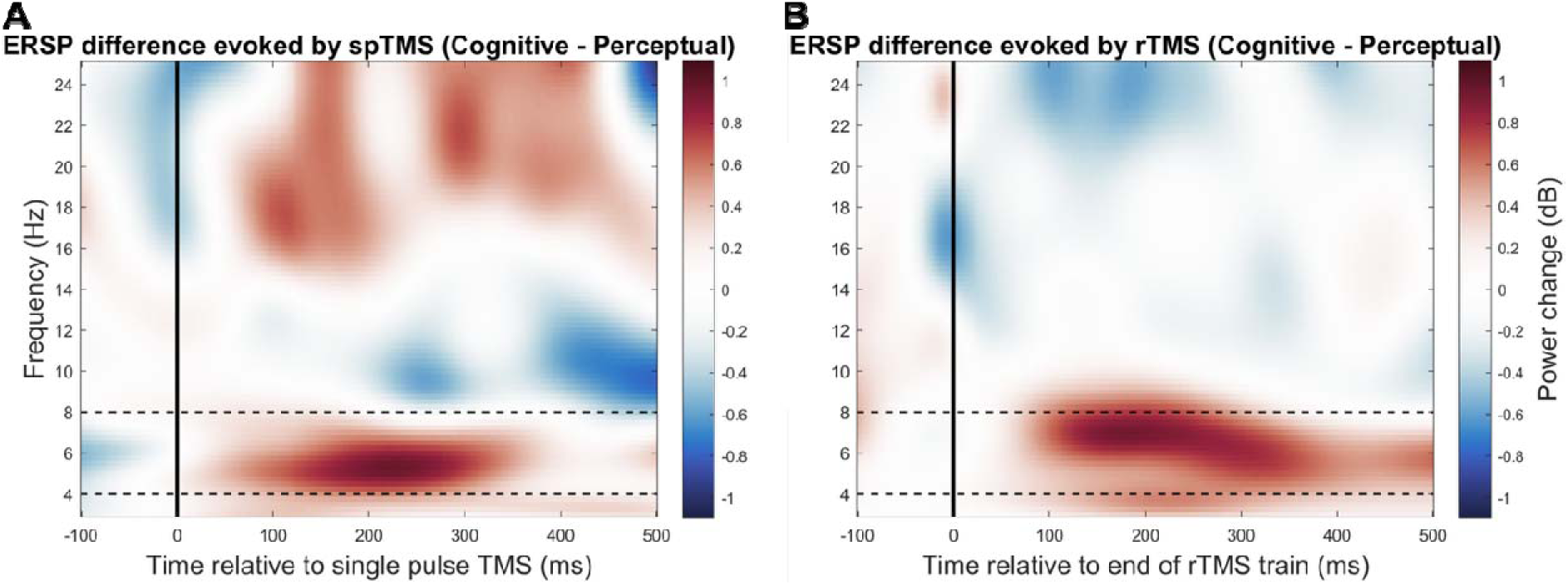
State-dependent theta band activation evoked by TMS. (A) Differences in ERSP power between TOPX and Perceptual tasks evoked by single-pulse TMS indicate an enhancement of TMS-evoked theta band oscillatory power during states of cognitive control relative to perceptual (control) states. (B) Similarly, trains of rTMS evoked stronger prefrontal oscillatory theta responses when applied during active behavioural states of cognitive control relative to perceptual states. Dashed lines represent the theta band spectral region for which we tested for condition differences via permutation tests. Power is computed against a divisive baseline in the linear power domain from -500 to -100 ms relative to the target event.

#### 3.2.2. State Dependent Modulation of Contextual EEG Responses

Cue-locked TOPX ERSPs were selectively modulated after TMS as a function of the behavioural state during TMS. Following ActiveState_rTMS_, the power of cue-locked theta-band ERSPs was significantly reduced relative to baseline, for both TOPX target cues, *t*_(22)_ = -4.13, *p* = .011, *d* = -0.95, and distractor (non-target) cues, *t*_(22)_ = -3.84, *p* = .039, *d* = -0.88. The effect was expressed over frontocentral and parietal electrodes for target cues, and left-prefrontal electrodes for distractor cues (**Figure 4B**). In contrast, when participants were engaged in the perceptual task during TMS, cluster-based tests showed no modulation of theta band activity for either TOPX target cues, *p* = .168, or distractor cues, *p* = .450. Moreover, there was spectral specificity in the state-dependent modulation of contextual neural processing. Significant time-frequency clusters of spatial cluster activity were limited to the theta and low-alpha bands for both TOPX target (latency = 38 – 786 ms, *t*_(22)_ = -7.36, *p* = .004, *d* = -1.69) and distractor (latency = 44 – 658 ms, *t*_(22)_ = -5.44, *p* = .022, *d* = -1.25) cues (**Figure 4C**).

**Figure 4.**
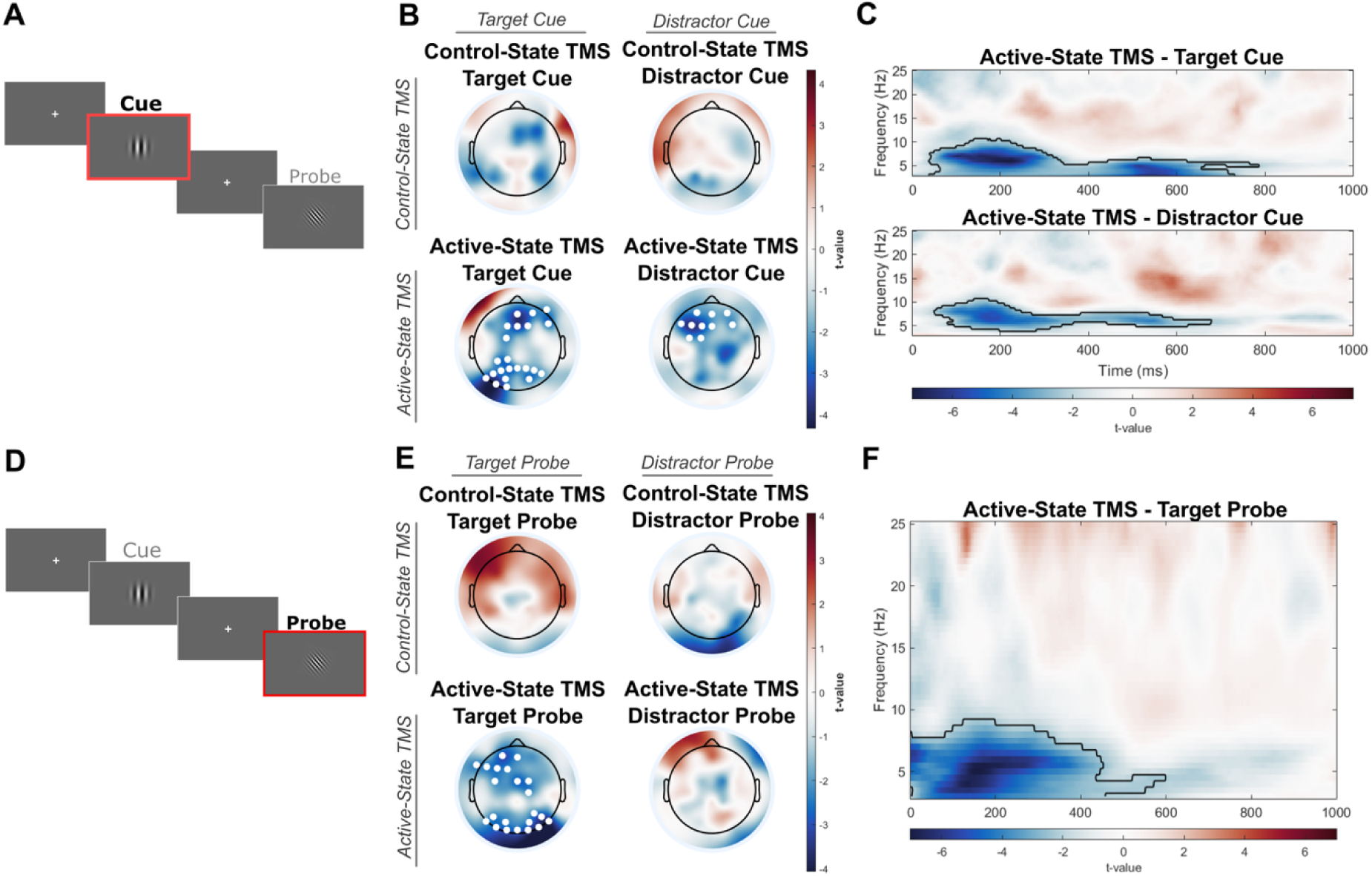
Modulation of goal maintenance and context processing by active-state TMS. (A) Theta-band oscillatory power evoked by TOPX cues was calculated as a difference from Pre- to Post- TMS, as a function of behavioural state at the time of TMS, showing a state-dependent modulation of context processing and goal maintenance by TMS. (B) The power of theta-band oscillatory responses was reduced after active-state TMS, but not control-state TMS. That modulation of theta-band activity showed a prefrontal-parietal distribution for target cues (consistent with reactive control), and a left-prefrontal distribution for distractor cues (consistent with proactive control). White circles show significant cluster-forming electrodes in cluster-based permutation tests. (C) The modulation of cue-evoked processing by active-state TMS was specific to theta and low-alpha bands. The outlined region shows significant time-frequency clusters in cluster-based permutation tests. (D) We also computed theta-band oscillatory power evoked by probe stimuli in the TOPX task, and calculated Pre- to Post- TMS differences in those ERSPs as a function of the behavioural state at the time of TMS. (E) The power of theta-band oscillatory responses to target probes was reduced after active-state TMS, showing a prefrontal-occipital distribution. White circles show significant cluster-forming electrodes in cluster-based permutation tests. (F) The modulation of target probe-evoked processing by active-state TMS was specific to theta and low-alpha bands. The outlined region shows significant time-frequency clusters in cluster-based permutation tests.

Probe-locked theta-band ERSPs were also reduced after ActiveState_rTMS_ for TOPX target probes, *t*_(22)_ = -3.87, *p* = .010, *d* = -0.89, but not distractor probes (*p* = .623). The target probe evoked cluster showed a left prefrontal-occipital distribution (**Figure 4E**) and spectral specificity to the theta and low-alpha bands (latency = 0 – 598 ms, *t*_(22)_ = -7.17, *p* = .010, *d* = -1.64; **Figure 4F**). In contrast, no significant clusters were detected for cue or distractor probe-locked ERSPs after ControlState_rTMS_.

Because sustained changes in pre-stimulus oscillatory activity could drive the observed state-dependent attenuation of theta ERSPs, we examined power spectral densities of the baseline periods used during computation of the ERSPs. Broadband spectral power increased on average after active-state TMS, but not control-state TMS (see **Supplementary Figure S3**).

However, at the level of individual subjects, those broadband power spectral shifts did not explain the magnitude of theta ERSP attenuation (see **Supplementary Materials** for a more detailed analysis). Therefore, the data suggest that TMS specifically modulated stimulus-locked theta activity in a state-dependent manner.

## 4. Discussion

Here, we demonstrated that the neuromodulatory effect of TMS on PFC-anchored networks is shaped by the cognitive state in which stimulation is delivered. When TMS was paired with active engagement of cognitive control circuits, participants showed selective enhancements of behavioural and neural indices that reflect the function of those circuits. In contrast, delivering the same stimulation during a perceptual discrimination task produced little measurable change. These findings provide direct evidence that endogenous engagement of PFC-anchored networks can sensitise those circuits to shape the magnitude and functional relevance of TMS-induced plasticity.

### 4.1. Modulation of prefrontal cortex circuitry by TMS is state dependent

Hebbian learning and metaplasticity models predict that active circuits are more receptive to modification^18,19^. Thus, actively engaging a target circuit in a desired state may bias the effects of neuromodulation. The notion that the ongoing state of neural activity can shape the propagation of TMS-evoked activity has been demonstrated across sensorimotor^16,27,29,75^, attention^30–33^, and memory^34–36^ domains. Here, we have shown the state dependence of TMS effects in complex cognitive function which spans across those domains. Relatedly, differential effects of TMS on affective processing have recently been observed to depend on trait-state depression severity at the time of stimulation^76^. Our results here highlight that states of cognition can be specifically manipulated to bias the effects of TMS toward clinically relevant outcomes. The effect on RT was smaller than we have previously reported with deep brain stimulation using a different cognitive control task^50^, but this might be expected for a non-invasive intervention.

### 4.2. Cognitive control as a mechanism for neural state manipulation

The finding that cognitive control can be selectively enhanced by TMS has direct clinical implications. Deficits in cognitive control are prevalent transdiagnostically^77–82^ and downstream readouts of that function can provide an objective measure of target circuit engagement^83,84^.

State-dependent circuit sensitisation might increase the clinical efficacy of neuromodulation and/or help target it to specific circuit deficits. Following active-state TMS, several downstream outputs of cognitive control were enhanced. Specifically, accuracy was higher, reflecting improved capacity for control, and probe RTs were shorter, suggesting enhanced computational efficiency. In addition, participants had a greater ability to use contextual information in support of goal-directed action – D’-Context was enhanced and decisional evidence was biased by sufficient contextual information (non-target cues). Moreover, we specifically showed in a follow-up experiment that practice/switching effects do not explain our results (see **Supplementary Materials**). An additional consideration is that we did not perform electric field modelling, which provides estimates of TMS-induced current distribution^85,86^. Importantly, such models do not specify which neural populations or networks are functionally engaged. Given that TMS-induced fields are relatively distributed across the cortex^87,88^, our study was not designed to rely on precise spatial localisation. Rather, we aimed to demonstrate that state-dependent circuit sensitisation plays a prominent role in stimulation outcomes. Together, our results suggest that TMS might be paired with structured cognitive exercises/training to increase the chance of response. Moreover, we observed an improvement of contextual processing, as manifested by D’-Context scores, but not in the index of proactive control (PBI_ER_, PBI_RT_). This suggests that active-state TMS did not push individuals into a more restrictive proactive/reactive regime, but enhanced the ability to flexibly adapt between modes of control. In other recent work, we showed that similar flexible switching may be a mechanism of recovery from depression^52^.

### 4.3. Mechanisms of cognitive control modulation

By using the context-dependent TOPX task to manipulate states of control, we leveraged a well-characterised mechanism for altering prefrontal activation and gating dynamics^63,89^. The two modes of control proposed by the DMC framework suggest flexible, context-dependent mechanisms that support goal directed behaviour. Proactive control actively sustains goal representations that bias attention and action systems^90^. Those biases are postulated to originate from anticipatory lateral PFC activity and a gating mechanism operating through corticostriatal circuits^89,91^. On the other hand, reactive control is transiently activated on an as-needed basis at the detection of conflict, triggering the retrieval and reorganisation of goal representations^90^. This process is thought to occur via a distinct and broader fronto-parietal network^91,92^. In addition, cognitive control is thought to operate through theta band activity which synchronises neural assemblies to integrate contextual information and enable flexible goal-directed control^37,71,92,93^. The spatial distribution and modulation within the theta band induced by ActiveState_rTMS_ is consistent with the above. Responses to non-target Distractor cues (which provide contextual information that permits proactive control) were modulated by active-state TMS, showing a reduction in theta band oscillatory power in a region over the left dorsolateral PFC. Moreover, we observed an increase in decisional evidence bias for non-target cues, which may reflect the effective engagement of proactive control. In contrast, target cues, for which reactive control may be required, evoked a fronto-parietal theta band modulation after active-state TMS. This suggests that both proactive and reactive control mechanisms may have been modified by active-state TMS.

We were surprised to observe a decrease, rather than an increase, in the power of task-evoked EEG responses after active-state TMS. For non-target cues, one interpretation is that this may reflect more active gating of PFC from subcortical regions, consistent with a proactive mode of control^89^. However, this does not explain the reduction of oscillatory theta power after (reactive) target cues and target probes. Given that behavioural metrics of control improved, a more parsimonious explanation is that reductions in theta band power may reflect an increase in the efficiency of the underlying computations. For example, prefrontal theta has been posed as a marker of the cognitive control demand or effort, rather than control ability^71,92,94^. Higher theta band oscillatory power may reflect compensatory recruitment of control resources when conflict is high or processing is inefficient. Thus, the attenuated theta band oscillatory power we observed, coupled with improved performance in downstream readouts of control, may suggest that active-state TMS was able to shift the system into a regime where less oscillatory drive was required to support goal-directed behaviour. This may reflect less effortful cognitive demands during conflict resolution, for instance, if prepotent responses are more easily inhibited after modulation of the underlying control networks. Additional peripheral measures of arousal associated with effort^95,96^ may help to delineate this further in the future.

### 4.4. Implications for understanding variability in the efficacy of TMS therapies

Our findings have implications for the mechanistic understanding of neuromodulation and its use clinically. Strategies to prime target circuits, such as engaging cognitive states, could improve treatment efficacy by increasing the physiologic receptivity of the target circuit. A limitation here is that our sample consisted of individuals without psychiatric disorders, to limit potential influence of heterogeneous or aberrant network processing. A key step forward would be to confirm that the same effects are observed in target populations, such as treatment resistant depression. Moreover, although improvements in cognitive control relate to clinical symptom improvement^50,52,97–100^, it remains to be tested whether the enhancement of control we observed can translate to clinically meaningful modifications of network function. Despite this, the modification of context processing by active-state TMS holds broad relevance as a therapeutic strategy across a range of disorders. Many conditions are characterised by impairments in maintaining or updating goal-relevant context, leading to inflexible or overly reactive thought and behaviour patterns. For example, impaired context sensitivity may contribute to persistent negative thinking and impaired expectation updating in individuals living with depression^101^.

Anxiety and phobias may involve exaggerated threat responses that persist even in the absence of contextual safety cues^102^. Similarly, obsessive-compulsive disorders may involve difficulty in using contextual information to suppress habitual behaviours^103^. As such, enhancing the function of circuits that support context processing and flexible control has broad potential to remediate a transdiagnostic marker of impairment^104^.

### 4.5. Conclusion

In summary, we have shown that the effects of prefrontal TMS are shaped by the ongoing state of cognition during stimulation. Actively engaging processes of cognitive control may sensitise underlying circuits to modification, which may bias the neuromodulatory effects to provide functional improvements in targeted neural circuits. Specifically, active-state TMS provided enhancements in downstream behavioural readouts and neural indices of PFC circuit function, which outlasted the period of stimulation. In contrast, TMS delivered during a non-active control state did not provide meaningful modification of those indices. This demonstration of state-dependent circuit sensitisation in prefrontal TMS can offer a path toward more reliable brain stimulation interventions.

## Acknowledgements

We thank the University of Minnesota Non-Invasive Neuromodulation Lab for their technical support and resources which supported this work.

## References

1. Blumberger DM, Mulsant BH, Daskalakis ZJ. What Is the Role of Brain Stimulation Therapies in the Treatment of Depression? Curr Psychiatry Rep. 2013;15(7):368. doi:10.1007/s11920-013-0368-1

2. Downar J, Siddiqi SH, Mitra A, Williams N, Liston C. Mechanisms of Action of TMS in the Treatment of Depression. In: Browning M, Cowen PJ, Sharp T, eds. Emerging Neurobiology of Antidepressant Treatments. Springer International Publishing; 2024:233–277. doi:10.1007/7854_2024_483

3. Hernández-Sauret A, Martin de la Torre O, Redolar-Ripoll D. Use of transcranial magnetic stimulation (TMS) for studying cognitive control in depressed patients: A systematic review. Cogn Affect Behav Neurosci. 2024;24(6):972–1007. doi:10.3758/s13415-024-01193-w

4. Chen L, Fukuda AM, Jiang S, et al. Treating Depression With Repetitive Transcranial Magnetic Stimulation: A Clinician’s Guide. Am J Psychiatry. Published online June 1, 2025. doi:10.1176/appi.ajp.20240859

5. Ge L, McInnes AN, Widge AS, Parhi KK. Determining the Number of Clusters in Clinical Response of TMS Treatment using Hyperdimensional Computing. J Signal Process Syst. Published online June 26, 2024. doi:10.1007/s11265-024-01921-y

6. Kaster TS, Downar J, Vila-Rodriguez F, et al. Differential symptom cluster responses to repetitive transcranial magnetic stimulation treatment in depression. eClinicalMedicine. 2023;55:101765. doi:10.1016/j.eclinm.2022.101765

7. McClintock SM, Reti IM, Carpenter LL, et al. Consensus recommendations for the clinical application of repetitive transcranial magnetic stimulation (rTMS) in the treatment of depression. J Clin Psychiatry. 2018;79(1):35–48. doi:10.4088/JCP.16cs10905

8. McInnes AN, Olsen ST, Sullivan CRP, et al. Trajectory modeling and response prediction in transcranial magnetic stimulation for depression. Pers Med Psychiatry. 2024;47–48:100135. doi:10.1016/j.pmip.2024.100135

9. Sackeim HA, Aaronson ST, Carpenter LL, et al. Clinical outcomes in a large registry of patients with major depressive disorder treated with Transcranial Magnetic Stimulation. J Affect Disord. 2020;277:65–74. doi:10.1016/j.jad.2020.08.005

10. Taylor SF, Bhati MT, Dubin MJ, et al. A naturalistic, multi-site study of repetitive transcranial magnetic stimulation therapy for depression. J Affect Disord. 2017;208:284–290. doi:10.1016/j.jad.2016.08.049

11. Bigoni C, Pagnamenta S, Cadic-Melchior A, et al. MEP and TEP features variability: is it just the brain-state? J Neural Eng. Published online 2024. doi:10.1088/1741-2552/ad1dc2

12. Bikson M, Rahman A. Origins of specificity during tDCS: anatomical, activity-selective, and input-bias mechanisms. Front Hum Neurosci. 2013;7. doi:10.3389/fnhum.2013.00688

13. Bradley C, Nydam AS, Dux PE, Mattingley JB. State-dependent effects of neural stimulation on brain function and cognition. Nat Rev Neurosci 2022 238. 2022;23(8):459-475. doi:10.1038/s41583-022-00598-1

14. Hussain SJ, Freedberg MV. Debunking the Myth of Excitatory and Inhibitory Repetitive Transcranial Magnetic Stimulation in Cognitive Neuroscience Research. J Cogn Neurosci. Published online December 22, 2024:1–14. doi:10.1162/jocn_a_02288

15. Sack AT, Paneva J, Küthe T, et al. Target engagement and brain state dependence of transcranial magnetic stimulation: implications for clinical practice. Biol Psychiatry. Published online September 21, 2023. doi:10.1016/j.biopsych.2023.09.011

16. Silvanto J, Muggleton N, Walsh V. State-dependency in brain stimulation studies of perception and cognition. Trends Cogn Sci. 2008;12(12):447–454. doi:10.1016/j.tics.2008.09.004

17. He H, Sun X, Doose J, et al. TMS-induced modulation of brain networks and its associations to rTMS treatment for depression: a concurrent fMRI-EEG-TMS study. Brain Stimulat. 2025;18(6):1955–1965. doi:10.1016/j.brs.2025.10.013

18. Hebb DO. The Organization of Behavior; a Neuropsychological Theory. Wiley; 1949:xix, 335.

19. Abraham WC. Metaplasticity: tuning synapses and networks for plasticity. Nat Rev Neurosci. 2008;9(5):387–387. doi:10.1038/nrn2356

20. Bloch J, Greaves-Tunnell A, Shea-Brown E, Harchaoui Z, Shojaie A, Yazdan-Shahmorad A. Network structure mediates functional reorganization induced by optogenetic stimulation of non-human primate sensorimotor cortex. iScience. 2022;25(5). doi:10.1016/j.isci.2022.104285

21. Fox MD, Buckner RL, White MP, Greicius MD, Pascual-Leone A. Efficacy of TMS targets for depression is related to intrinsic functional connectivity with the subgenual cingulate. Biol Psychiatry. 2012;72(7):595–603. doi:10.1016/j.biopsych.2012.04.028

22. Neacsiu A, Luber BM, Davis S, Bernhardt E, Strauman TJ, Lisanby SH. On the Concurrent Use of Self-System Therapy (SST) and fMRI-guided Transcranial Magnetic Stimulation (TMS) as Treatment for Depression. J ECT. 2018;34(4):266–273. doi:10.1097/YCT.0000000000000545

23. Xu X, Xu M, Su Y, et al. Efficacy of Repetitive Transcranial Magnetic Stimulation (rTMS) Combined with Psychological Interventions: A Systematic Review and Meta-Analysis of Randomized Controlled Trials. Brain Sci. 2023;13(12):1665. doi:10.3390/brainsci13121665

24. Conelea CA, Jacob S, Redish AD, Ramsay IS. Considerations for Pairing Cognitive Behavioral Therapies and Non-invasive Brain Stimulation: Ignore at Your Own Risk. Front Psychiatry. 2021;12. doi:10.3389/fpsyt.2021.660180

25. Carmi L, Tendler A, Bystritsky A, et al. Efficacy and Safety of Deep Transcranial Magnetic Stimulation for Obsessive-Compulsive Disorder: A Prospective Multicenter Randomized Double-Blind Placebo-Controlled Trial. Am J Psychiatry. 2019;176(11):931–938. doi:10.1176/appi.ajp.2019.18101180

26. Cavallero F, Gold MC, Tirrell E, et al. Audio-Guided Mindfulness Meditation During Transcranial Magnetic Stimulation Sessions for the Treatment of Major Depressive Disorder: A Pilot Feasibility Study. Front Psychol. 2021;12. doi:10.3389/fpsyg.2021.678911

27. Decroix J, Borgomaneri S, Kalénine S, Avenanti A. State-dependent TMS of inferior frontal and parietal cortices highlights integration of grip configuration and functional goals during action recognition. Cortex. 2020;132:51–62. doi:10.1016/j.cortex.2020.08.004

28. McInnes AN, Smithers B, Lipp OV, et al. From Inhibition to Excitation and Why: The Role of Temporal Urgency in Modulating Corticospinal Activity. bioRxiv. Preprint posted online December 28, 2023:2023.12.27.573452. doi:10.1101/2023.12.27.573452

29. Silvanto J, Muggleton NG, Cowey A, Walsh V. Neural adaptation reveals state-dependent effects of transcranial magnetic stimulation. Eur J Neurosci. 2007;25(6):1874–1881. doi:10.1111/j.1460-9568.2007.05440.x

30. Capotosto P, Baldassarre A, Sestieri C, Spadone S, Romani GL, Corbetta M. Task and Regions Specific Top-Down Modulation of Alpha Rhythms in Parietal Cortex. Cereb Cortex. 2017;27(10):4815–4822. doi:10.1093/cercor/bhw278

31. Heinen K, Feredoes E, Weiskopf N, Ruff CC, Driver J. Direct Evidence for Attention-Dependent Influences of the Frontal Eye-Fields on Feature-Responsive Visual Cortex. Cereb Cortex. 2014;24(11):2815–2821. doi:10.1093/cercor/bht157

32. Herring JD, Thut G, Jensen O, Bergmann TO. Attention Modulates TMS-Locked Alpha Oscillations in the Visual Cortex. J Neurosci. 2015;35(43):14435–14447. doi:10.1523/JNEUROSCI.1833-15.2015

33. Kamke MR, Ryan AE, Sale MV, et al. Visual Spatial Attention Has Opposite Effects on Bidirectional Plasticity in the Human Motor Cortex. J Neurosci. 2014;34(4):1475–1480. doi:10.1523/JNEUROSCI.1595-13.2014

34. Albouy P, Weiss A, Baillet S, Zatorre RJ. Selective Entrainment of Theta Oscillations in the Dorsal Stream Causally Enhances Auditory Working Memory Performance. Neuron. 2017;94(1):193–206.e5. doi:10.1016/j.neuron.2017.03.015

35. Feredoes E, Heinen K, Weiskopf N, Ruff C, Driver J. Causal evidence for frontal involvement in memory target maintenance by posterior brain areas during distracter interference of visual working memory. Proc Natl Acad Sci. 2011;108(42):17510–17515. doi:10.1073/pnas.1106439108

36. Rose NS, LaRocque JJ, Riggall AC, et al. Reactivation of latent working memories with transcranial magnetic stimulation. Science. 2016;354(6316):1136–1139. doi:10.1126/science.aah7011

37. Menon V, D’Esposito M. The role of PFC networks in cognitive control and executive function. Neuropsychopharmacology. 2022;47(1):90–103. doi:10.1038/s41386-021-01152-w

38. Dengler J, Deck BL, Stoll H, et al. Enhancing cognitive control with transcranial magnetic stimulation in subject-specific frontoparietal networks. Cortex. 2024;172:141–158. doi:10.1016/j.cortex.2023.11.020

39. Sachse EM, Widge AS. Neurostimulation to improve cognitive flexibility. Curr Opin Behav Sci. 2025;62:101484. doi:10.1016/j.cobeha.2025.101484

40. Rossi S, Antal A, Bestmann S, et al. Safety and recommendations for TMS use in healthy subjects and patient populations, with updates on training, ethical and regulatory issues: Expert Guidelines. Clin Neurophysiol. 2021;132(1):269–306. doi:10.1016/J.CLINPH.2020.10.003

41. Green P, MacLeod CJ, Nakagawa S. SIMR: an R package for power analysis of generalized linear mixed models by simulation. Methods Ecol Evol. 2016;7(4):493–498. doi:10.1111/2041-210X.12504

42. Brysbaert M, Stevens M. Power Analysis and Effect Size in Mixed Effects Models: A Tutorial. J Cogn. 2018;1(1):9. doi:10.5334/joc.10

43. Brunoni AR, Vanderhasselt MA. Working memory improvement with non-invasive brain stimulation of the dorsolateral prefrontal cortex: A systematic review and meta-analysis. Brain Cogn. 2014;86:1–9. doi:10.1016/j.bandc.2014.01.008

44. Beynel L, Deng L, Crowell CA, et al. Structural Controllability Predicts Functional Patterns and Brain Stimulation Benefits Associated with Working Memory. J Neurosci. 2020;40(35):6770–6778. doi:10.1523/JNEUROSCI.0531-20.2020

45. Seeck M, Koessler L, Bast T, et al. The standardized EEG electrode array of the IFCN. Clin Neurophysiol. 2017;128(10):2070–2077. doi:10.1016/j.clinph.2017.06.254

46. Arend JL. Observed and Drift Diffusion Modeled Performance in Early Psychosis: Decreased Drift Rates and Bias in a Cognitive Control Task. M.A. University of Minnesota; 2024. Accessed November 24, 2025. https://www.proquest.com/docview/3082376592/abstract/EE6947E50B26468BPQ/1

47. R Core Team. A language and environment for statistical computing. R Foundation for Statistical Computing, Vienna (2016). Published online 2016:2013.

48. Basu I, Yousefi A, Crocker B, et al. Closed-loop enhancement and neural decoding of cognitive control in humans. Nat Biomed Eng. 2023;7(4):576–588. doi:10.1038/s41551-021-00804-y

49. Reimer AE, Dastin-van Rijn EM, Kim J, et al. Striatal stimulation enhances cognitive control and evidence processing in rodents and humans. Sci Transl Med. 2024;16(778):eadp1723. doi:10.1126/scitranslmed.adp1723

50. Widge AS, Zorowitz S, Basu I, et al. Deep brain stimulation of the internal capsule enhances human cognitive control and prefrontal cortex function. Nat Commun. 2019;10(1):1536. doi:10.1038/s41467-019-09557-4

51. Glewwe N, Rijn EDV, Chen CS, et al. Sex-biased computations underlying differential set shift performance in mice. bioRxiv. Preprint posted online April 3, 2025:2025.04.01.646712. doi:10.1101/2025.04.01.646712

52. Kim J, Widge AS. Cingulate-centered flexible control: physiologic correlates and enhancement by internal capsule stimulation. bioRxiv. Preprint posted online October 16, 2025:2025.10.15.682151. doi:10.1101/2025.10.15.682151

53. Sachse EM, Rijn EMD van, Bennek JP, et al. Unilateral striatal deep brain stimulation improves cognitive control. bioRxiv. Preprint posted online November 12, 2025:2025.11.10.686309. doi:10.1101/2025.11.10.686309

54. Cole RC, Ging-Jehli NR, Vivanco Suarez J, et al. Theta-frequency subthalamic nucleus stimulation increases decision threshold. Brain Stimulat. 2025;18(4):1021–1027. doi:10.1016/j.brs.2025.05.105

55. Georgiev D, Rocchi L, Tocco P, Speekenbrink M, Rothwell JC, Jahanshahi M. Continuous Theta Burst Stimulation Over the Dorsolateral Prefrontal Cortex and the Pre-SMA Alter Drift Rate and Response Thresholds Respectively During Perceptual Decision-Making. Brain Stimulat. 2016;9(4):601–608. doi:10.1016/j.brs.2016.04.004

56. Liu X, Guo T, Chang Q, et al. Optimizing cognitive control through the interaction between stimulation intensity and duration in single-site and dual-site tDCS. Sci Rep. 2025;15(1):35963. doi:10.1038/s41598-025-14509-8

57. Shen C, Calvin OL, Rawls E, Redish AD, Sponheim SR. Clarifying Cognitive Control Deficits in Psychosis via Drift Diffusion Modeling and Attractor Dynamics. Schizophr Bull. Published online February 26, 2024:sbae014. doi:10.1093/schbul/sbae014

58. Delorme A, Makeig S. EEGLAB: an open source toolbox for analysis of single-trial EEG dynamics including independent component analysis. J Neurosci Methods. 2004;134(1):9–21. doi:10.1016/j.jneumeth.2003.10.009

59. Oostenveld R, Fries P, Maris E, Schoffelen JM. FieldTrip: Open Source Software for Advanced Analysis of MEG, EEG, and Invasive Electrophysiological Data. Comput Intell Neurosci. 2011;2011:1–9. doi:10.1155/2011/156869

60. Rogasch NC, Sullivan C, Thomson RH, et al. Analysing concurrent transcranial magnetic stimulation and electroencephalographic data: A review and introduction to the open-source TESA software. NeuroImage. 2017;147:934–951. doi:10.1016/j.neuroimage.2016.10.031

61. Rogasch NC, Thomson RH, Farzan F, et al. Removing artefacts from TMS-EEG recordings using independent component analysis: importance for assessing prefrontal and motor cortex network properties. NeuroImage. 2014;101:425–439. doi:10.1016/j.neuroimage.2014.07.037

62. Tallon-Baudry C, Bertrand O. Oscillatory gamma activity in humans and its role in object representation. Trends Cogn Sci. 1999;3(4):151–162. doi:10.1016/S1364-6613(99)01299-1

63. Braver TS, Paxton JL, Locke HS, Barch DM. Flexible neural mechanisms of cognitive control within human prefrontal cortex. Proc Natl Acad Sci. 2009;106(18):7351–7356. doi:10.1073/PNAS.0808187106

64. Barch DM, Berman MG, Engle R, et al. CNTRICS Final Task Selection: Working Memory. Schizophr Bull. 2009;35(1):136–152. doi:10.1093/schbul/sbn153

65. Cohen JD, Servan-Schreiber D. Context, cortex, and dopamine: A connectionist approach to behavior and biology in schizophrenia. Psychol Rev. 1992;99(1):45–77. doi:10.1037/0033-295X.99.1.45

66. Cooper SR, Gonthier C, Barch DM, Braver TS. The Role of Psychometrics in Individual Differences Research in Cognition: A Case Study of the AX-CPT. Front Psychol. 2017;8. doi:10.3389/fpsyg.2017.01482

67. MacDonald III AW, Goghari VM, Hicks BM, Flory JD, Carter CS, Manuck SB. A Convergent-Divergent Approach to Context Processing, General Intellectual Functioning, and the Genetic Liability to Schizophrenia. Neuropsychology. 2005;19(6):814–821. doi:10.1037/0894-4105.19.6.814

68. Servan-Schreiber D, Cohen JD, Steingard S. Schizophrenic Deficits in the Processing of Context: A Test of a Theoretical Model. Arch Gen Psychiatry. 1996;53(12):1105–1112. doi:10.1001/archpsyc.1996.01830120037008

69. Braver TS. The variable nature of cognitive control: a dual mechanisms framework. Trends Cogn Sci. 2012;16(2):106–113. doi:10.1016/j.tics.2011.12.010

70. Ahumada-Méndez F, Lucero B, Avenanti A, et al. Affective modulation of cognitive control: A systematic review of EEG studies. Physiol Behav. 2022;249:113743. doi:10.1016/j.physbeh.2022.113743

71. Cavanagh JF, Frank MJ. Frontal theta as a mechanism for cognitive control. Trends Cogn Sci. 2014;18(8):414–421. doi:10.1016/j.tics.2014.04.012

72. Li Q, Chen T, Wang L, et al. Novelty modulates proactive and reactive cognitive control modes: Evidence from ERP and EEG data. NeuroImage. 2025;311:121178. doi:10.1016/j.neuroimage.2025.121178

73. Soltani Zangbar H, Ghadiri T, Seyedi Vafaee M, et al. Theta Oscillations Through Hippocampal/Prefrontal Pathway: Importance in Cognitive Performances. Brain Connect. 2020;10(4):157–169. doi:10.1089/brain.2019.0733

74. Gordon PC, Belardinelli P, Stenroos M, Ziemann U, Zrenner C. Prefrontal theta phase-dependent rTMS-induced plasticity of cortical and behavioral responses in human cortex. Brain Stimulat. 2022;15(2):391–402. doi:10.1016/j.brs.2022.02.006

75. McInnes AN, Lipp OV, Tresilian JR, Vallence AM, Marinovic W. Premovement inhibition can protect motor actions from interference by response-irrelevant sensory stimulation. J Physiol. 2021;599(18):4389–4406. doi:10.1113/JP281849

76. Paneva J, Schuhmann T, De Smet S, De Meza T, Duecker F, Sack AT. Affective state-dependent effects of prefrontal rTMS on the cognitive control of negative stimuli in healthy and depressed individuals. Brain Stimulat. 2025;18(3):745–752. doi:10.1016/j.brs.2025.04.002

77. Breukelaar IA, Erlinger M, Harris A, et al. Investigating the neural basis of cognitive control dysfunction in mood disorders. Bipolar Disord. 2020;22(3):286–295. doi:10.1111/bdi.12844

78. Dalley JW, Everitt BJ, Robbins TW. Impulsivity, Compulsivity, and Top-Down Cognitive Control. Neuron. 2011;69(4):680–694. doi:10.1016/j.neuron.2011.01.020

79. McInnes AN, Sullivan CRP, Angus W. MacDonald III, Widge AS. Psychometric Validation and Preliminary Clinical Correlation of an Experiential Foraging Task. Assessment. Published online September 25, 2025. doi:10.1177/10731911251376214

80. McTeague LM, Goodkind MS, Etkin A. Transdiagnostic impairment of cognitive control in mental illness. J Psychiatr Res. 2016;83:37–46. doi:10.1016/j.jpsychires.2016.08.001

81. McTeague LM, Huemer J, Carreon DM, Jiang Y, Eickhoff SB, Etkin A. Identification of Common Neural Circuit Disruptions in Cognitive Control Across Psychiatric Disorders. Am J Psychiatry. 2017;174(7):676–685. doi:10.1176/appi.ajp.2017.16040400

82. Widge AS. Closing the loop in psychiatric deep brain stimulation: physiology, psychometrics, and plasticity. Neuropsychopharmacol Off Publ Am Coll Neuropsychopharmacol. 2024;49(1):138–149. doi:10.1038/s41386-023-01643-y

83. Nagrale SS, Yousefi A, Netoff TI, Widge AS. In silico development and validation of Bayesian methods for optimizing deep brain stimulation to enhance cognitive control. J Neural Eng. 2023;20(3):036015. doi:10.1088/1741-2552/acd0d5

84. Rijn EMD van, Sachse EM, Buccini M, et al. Real-time Bayesian optimization of deep brain stimulation for personalized cognitive control enhancement. bioRxiv. Preprint posted online December 30, 2025:2025.12.30.697057. doi:10.64898/2025.12.30.697057

85. Opitz A, Windhoff M, Heidemann RM, Turner R, Thielscher A. How the brain tissue shapes the electric field induced by transcranial magnetic stimulation. Neuroimage. 2011;58(3):849–859.

86. Dannhauer M, Gomez LJ, Robins PL, et al. Electric field modeling in personalizing TMS interventions. Biol Psychiatry. 2024;95(6):494–501. doi:10.1016/j.biopsych.2023.11.022

87. Tadayon P, MacKenzie A, Gregory E, Pichardo S, Görges M, Vila-Rodriguez F. Quantification of Transcranial Magnetic Stimulation and Low-intensity Focused Ultrasound Energy Field Focality in the Cerebral Cortex. Neuromodulation Technol Neural Interface. Published online February 27, 2026. doi:10.1016/j.neurom.2026.01.014

88. Siebner HR, Funke K, Aberra AS, et al. Transcranial magnetic stimulation of the brain: What is stimulated? – A consensus and critical position paper. Clin Neurophysiol Off J Int Fed Clin Neurophysiol. 2022;140:59–97. doi:10.1016/j.clinph.2022.04.022

89. Chiew KS, Braver TS. Context Processing and Cognitive Control: From Gating Models to Dual Mechanisms. In: Egner T, ed. The Wiley Handbook of Cognitive Control. 1st ed. Wiley; 2017:143-166. doi:10.1002/9781118920497.ch9

90. Etzel JA, Brough RE, Freund MC, et al. The Dual Mechanisms of Cognitive Control dataset, a theoretically-guided within-subject task fMRI battery. Sci Data. 2022;9(1):114. doi:10.1038/s41597-022-01226-4

91. Kwashie AND, Ma Y, Barch DM, et al. Comparing the Functional Neuroanatomy of Proactive and Reactive Control between Patients with Schizophrenia and Healthy Controls. Cogn Affect Behav Neurosci. 2023;23(1):203–215. doi:10.3758/s13415-022-01036-6

92. Cooper PS, Wong ASW, Fulham WR, et al. Theta frontoparietal connectivity associated with proactive and reactive cognitive control processes. NeuroImage. 2015;108:354–363. doi:10.1016/j.neuroimage.2014.12.028

93. Riddle J, Vogelsang DA, Hwang K, Cellier D, D’Esposito M. Distinct Oscillatory Dynamics Underlie Different Components of Hierarchical Cognitive Control. J Neurosci. 2020;40(25):4945–4953. doi:10.1523/JNEUROSCI.0617-20.2020

94. McFerren A, Riddle J, Walker C, Buse JB, Frohlich F. Causal role of frontal-midline theta in cognitive effort: a pilot study. J Neurophysiol. 2021;126(4):1221–1233. doi:10.1152/jn.00068.2021

95. Kuipers M, Richter M, Scheepers D, Immink MA, Sjak-Shie E, van Steenbergen H. How effortful is cognitive control? Insights from a novel method measuring single-trial evoked beta-adrenergic cardiac reactivity. Int J Psychophysiol. 2017;119:87–92. doi:10.1016/j.ijpsycho.2016.10.007

96. McInnes AN, Sung B. A neglected consumer neuroscience technique: Pupillometry and its practical application to consumer research. Int J Res Mark. 2025;42(3, Part B):827–843. doi:10.1016/j.ijresmar.2024.11.005

97. Crane NA, Jenkins LM, Bhaumik R, et al. Multidimensional prediction of treatment response to antidepressants with cognitive control and functional MRI. Brain. 2017;140(2):472–486. doi:10.1093/brain/aww326

98. Ge L, McInnes AN, Widge AS, Parhi KK. Prediction of Clinical Response of Transcranial Magnetic Stimulation Treatment for Major Depressive Disorder Using Hyperdimensional Computing. IEEE J Biomed Health Inform. Published online 2025:1–9. doi:10.1109/JBHI.2025.3537757

99. Hack LM, Tozzi L, Zenteno S, et al. A Cognitive Biotype of Depression Linking Symptoms, Behavior Measures, Neural Circuits, and Differential Treatment Outcomes: A Prespecified Secondary Analysis of a Randomized Clinical Trial. JAMA Netw Open. 2023;6(6):e2318411. doi:10.1001/jamanetworkopen.2023.18411

100. Koster EHW, Hoorelbeke K, Onraedt T, Owens M, Derakshan N. Cognitive control interventions for depression: A systematic review of findings from training studies. Clin Psychol Rev. 2017;53:79–92. doi:10.1016/j.cpr.2017.02.002

101. Stange JP, Connolly SL, Burke TA, et al. Inflexible Cognition Predicts First Onset of Major Depressive Episodes in Adolescence. Depress Anxiety. 2016;33(11):1005–1012. doi:10.1002/da.22513

102. Abend R. Understanding anxiety symptoms as aberrant defensive responding along the threat imminence continuum. Neurosci Biobehav Rev. 2023;152:105305. doi:10.1016/j.neubiorev.2023.105305

103. Weiss F, Schwarz K, Endrass T. Exploring the relationship between context and obsessions in individuals with obsessive-compulsive disorder symptoms: a narrative review. Front Psychiatry. 2024;15:1353962. doi:10.3389/fpsyt.2024.1353962

104. Tozzi L, Bertrand C, Hack LM, et al. A cognitive neural circuit biotype of depression showing functional and behavioral improvement after transcranial magnetic stimulation in the B-SMART-fMRI trial. Nat Ment Health. 2024;2(8):987–998. doi:10.1038/s44220-024-00271-9

