## Supplementary Materials for "Cognitive state dependent enhancement of cognitive control with transcranial magnetic stimulation"

Single pulse and repetitive TMS was applied using a Magstim Super Rapid$^2$ Plus stimulator connected to a figure-of-eight Air Film Coil with a wing diameter of 70 mm (Magstim Co. Ltd, Wales, UK). TMS was applied at 100% of the participant's RMT (to the nearest 1% of maximum stimulator output) over the left PFC using the Beam-F3 targeting method (Beam et al., 2009). We applied TMS conservatively at 100% of RMT, rather than the conventional intensity of 120%, for two reasons. First, stimulation near threshold is more likely to preserve sensitivity to ongoing neural state, which was central to our hypothesis regarding state-dependent effects of TMS. Second, using a lower intensity reduced the risk of coil overheating, particularly in participants with higher resting motor thresholds (RMTs).

Prior to beginning experimental trials, we first identified the scalp location of the first dorsal interosseous (FDI) hotspot, and determined each participant's RMT using the Rossini-Rothwell procedure (Rossini et al., 1994). This procedure involved applying single-pulse TMS over the right FDI hotspot with the muscle at rest to identify the minimum stimulation intensity (in % of maximum stimulator output) at which TMS could elicit a motor evoked potential with an amplitude greater than 50 µV in at least 5 out of 10 trials. We recorded the amplitude of those motor evoked potentials using an electromyography (EMG) amplifier (EMG Pod, Rogue Research Inc., Canada) with surface 8 mm Ag/AgCl sintered electrodes over the right FDI in a monopolar muscle belly-tendon montage and with a reference electrode placed over the styloid process of the right ulna. The mean resting motor threshold was 65.95% of maximum stimulator output (SD = 7.35, range = 51% - 86%). Resting motor thresholds showed excellent consistency within subjects across both testing sessions, with an intraclass correlation coefficient = 0.91, $p < .001$. Repeated testing sessions were conducted at approximately the same time of day within participants to limit circadian influences on cortical excitability (Ly et al., 2016).

Throughout all TMS procedures, the stimulation coil was placed tangentially to the skull with the handle rotated clockwise 45 degrees from the midsagittal plane. Consistency in the position and orientation of the coil was maintained throughout the experiment using frameless stereotactic neuronavigation (BrainSight *v2.5.8,* Rogue Research Inc., Canada). Stimulation parameters were controlled via MATLAB using the MAGIC toolbox (*v1.0*, Habibollahi Saatlou et al., 2018). To reduce auditory evoked potentials from the coil's discharge (Biabani et al., 2024), participants wore hearing protective earplugs, and active noise-cancelling headphones (BOSE QC25) playing white noise at an amplitude of approximately 75 dBA throughout the experiment, including task blocks where no stimulation was applied.

During rTMS blocks, trains of rTMS were delivered during the inter-trial interval in a modified intermittent burst stimulation pattern (**Figure 1**). Stimulation was applied during the inter-trial interval to avoid facial and ocular muscle contraction during stimulus presentation that may have interfered with task performance. Bursts of three TMS pulses with an inter-pulse interval of 20 ms (50 Hz) were triggered every 333 ms (3 Hz) and 10 bursts were applied every trial. Trial durations resulted in an inter-train interval ranging from 9.5 – 11.5 s. The carrier frequency was slower than that of traditional intermittent theta burst stimulation, due to limitations of the stimulator in triggering bursts at shorter intervals when parameters were set externally via the MAGIC toolbox. We chose to apply an intermittent burst-like pattern of stimulation because it provided a natural temporal alignment with the task inter-trial interval, which allowed us to deliver stimulation intermittently between trials without overlap with task performance. In a subset (26.66%) of trials in both the $\text{ActiveState}_{\text{rTMS}}$ and $\text{ControlState}_{\text{rTMS}}$ tasks, we delivered single-pulse TMS during the inter-stimulus interval to probe cortical excitability and responsiveness to stimulation. During $\text{ActiveState}_{\text{rTMS}}$ blocks, single-pulse TMS was applied in 16 (22%) of AX trials, and in 8 trials (44%) each of the BX and BY trials (refer to **Figure 1** for trial types in the TOPX task). In the $\text{ControlState}_{\text{rTMS}}$ condition, single pulse TMS was applied in an equal subset of trials presenting the cue-probe stimuli in Vertical-Vertical, Vertical-Horizontal, Horizontal-Vertical, and Horizontal-Horizontal orientations. Trial distributions were pseudorandom so that single pulse TMS was not presented in any two consecutive trials.

*Cognitive control and perceptual behavioural tasks*

TOPX builds upon the AX-CPT (Cohen et al., 1997; Cohen & Servan-Schreiber, 1992; Servan-Schreiber et al., 1996) framework, which assesses context processing and goal maintenance. The task presents two visual stimuli, a cue and a probe, each requiring a behavioural response. A response is given at presentation of the target cue-probe sequence, and an alternate response is required for non-target cue-probe sequences (refer to **Figure 1**). The target sequence is presented in a majority of trials to produce a response bias that must be countermanded in non-target trials. Due to this response bias, following target cues, there is an expectancy that the subsequent probe trial will require a target response. This target expectancy produces interference when a non-target probe is presented instead, requiring cognitive control to resolve that conflict. Similarly, when a non-target cue is presented, participants must overcome the target response bias if that non-target cue is followed by a target probe.

The perceptual discrimination task had the same stimulus parameters as TOPX, but did not require context processing or inhibition of prepotent responses. Rather than responding to target/distractor stimulus sequences, participants responded according to the orientation of each stimulus that was displayed. Correct responses required pressing the down arrow key for vertically oriented Gabor patches and the left arrow key for horizontally oriented Gabor patches. In each trial, two patches were presented, each with a 50% probability of vertical/horizontal orientation. Participants were given feedback of a correct response if they correctly identified the orientation of both Gabor patches in a trial (matching the two-stimulus cue-probe sequence of TOPX). The experimental tasks were administered in MATLAB using Psychtoolbox (Kleiner et al., 2007).

*Behavioural data processing and analysis*

We screened behavioural data for blocks where response accuracy was < 70% or where the number of omitted responses was > 10. No blocks met these criteria. Two participants did not complete the second experimental session and were excluded from analysis. To assess cognitive control performance before and after stimulation, we computed accuracy, probe reaction times (RTs), D′-Context, and the Proactive Behavioural Index of error rates ($PBI_{ER}$) and probe RTs ($PBI_{RT}$) for each run of TOPX. Accuracy was computed as the number of correct responses divided by the number of trials in each run. RTs were registered by the task as the time difference between stimulus onset and the time a valid keypress was detected. We removed probe RTs < 200 ms (indicating anticipatory responses; 0.52%) and RTs > 1500 ms (indicating lapsed attention; 0.76%) from analysis. D′-Context was calculated as a measure of context processing, as the z-score difference between the proportion of correct responses in the AX trial sequence (hit rate) and the proportion of incorrect responses in the BX trial sequence (false alarm rate); $D'\text{-}Context = z(AX_{Hit}) - z(BX_{FalseAlarm})$ (Servan-Schreiber et al., 1996). $PBI_{ER}$ and $PBI_{RT}$ were calculated as a measure of proactive control over reactive control, as the ratio of errors and reaction times, respectively, in proactive versus reactive trial types; *PBI = (AY – BX)/(AY + BX)* (Braver et al., 2009).

We assessed change in TOPX accuracy, D′-Context, and $PBI_{ER}$ across blocks (Pre/Post) as a function of behavioural state at the time of TMS ($ActiveState_{rTMS}$, $ControlState_{rTMS}$) using a series of linear mixed-effects models, including session number (1, 2) as a main effect and subject IDs as a random intercept. Given their positive, right-skewed distribution, we analysed RTs using a generalised linear mixed effects model with a gamma link function and identity distribution, as in our past work (Reimer et al., 2024; Widge et al., 2019), applying the same fixed effects and random intercept structure as the linear mixed model. We report $R^2$ values alongside our statistical tests as an estimate of effect size. Linear models were run with the Satterthwaite approximation for degrees of freedom (Satterthwaite, 1946) and post-hoc tests were corrected for multiple comparisons using the false discovery rate method (Benjamini & Hochberg, 1995).

We also fit hierarchical drift diffusion models (DDMs) to reaction times and response accuracy to infer latent processes of evidence accumulation towards decisions in TOPX. For each run of TOPX, we estimated the drift rate ($v$), decision threshold ($a$), non-decision time ($t$), and starting point bias ($z$) using the HDDM package (Wiecki et al., 2013). Consistent with a previous report applying hierarchical DDMs in the dot-pattern expectancy task (Shen et al., 2024), a paradigm similar to TOPX, we found the best fitting model according to posterior predictive checks and Deviance Information Criterion (DIC) permitted the $v$ and $a$ parameters to vary by trial sequence (AX, AY, BX, BY), while $z$ was permitted to vary only by cue type (A Cue, B Cue), and t was held constant across conditions and estimated hierarchically with a group-level prior. This model yielded a mean DIC across chains = -19307.31, and a ΔDIC = -73.2 relative to the next best converging model which permitted only $z$ to vary by trial sequence. We ran five independent chains per model, and we assessed model convergence by confirming that the Gelman-Rubin $\hat{R}$ (Gelman & Rubin, 1992) statistic was < 1.1.

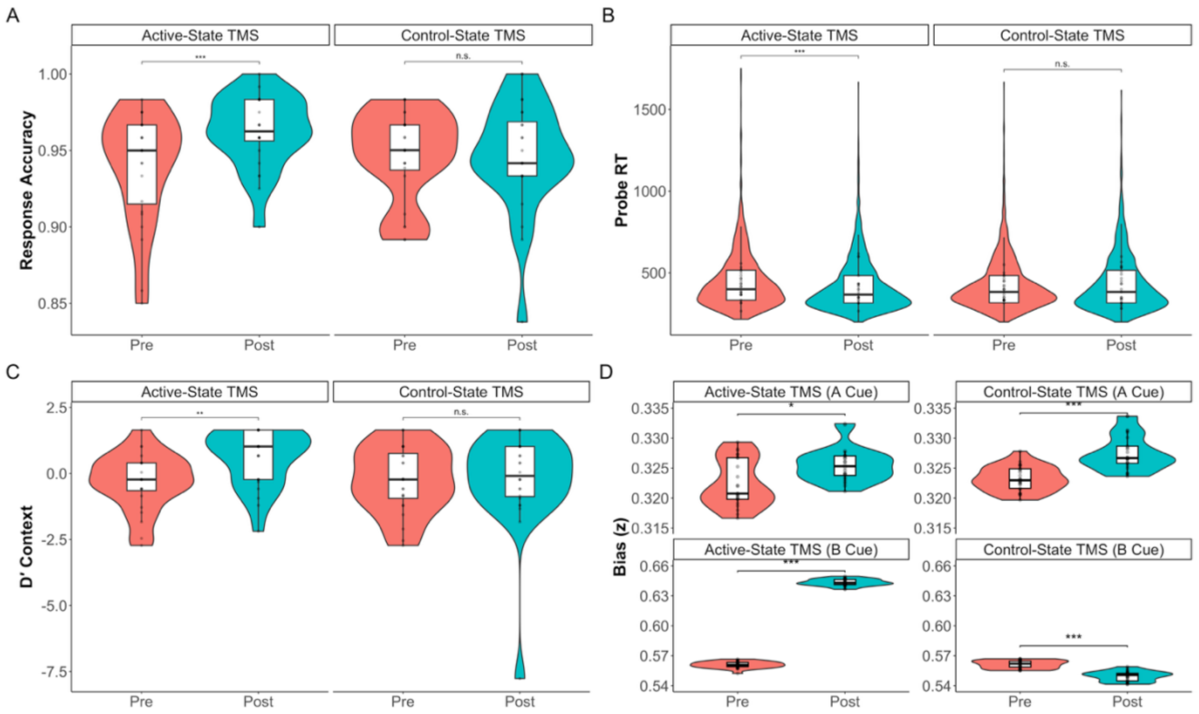


**Figure S1**. TOPX raw pre- and post-rTMS behavioural results in Experiment One as a function of the behavioural state at the time of stimulation. (A) After active-state TMS, response accuracy increased, but not after control-state TMS. (B) Probe reaction times (RTs) shortened after active-state, but not control-state TMS. (C) Contextual discrimination was also selectively enhanced after active-state TMS. (D) Evidence accumulation bias estimated by drift diffusion modelling showed a state dependent modulation by TMS, particularly for non-target cues, whereby starting point bias was increased in active-state TMS, but decreased in the control-state TMS condition. Violins depict probability density distributions for each condition; boxplots depict the between-subject median and interquartile range for each condition; points represent individual-subject change scores. For probe RTs (B), violins, boxplots, and statistical significance are depicted for trial-level distributions, and points represent medians computed for each subject. For all other behavioural metrics, violins, boxplots, points, and significance represent the distribution of means calculated for each participant. Significance (FDR-corrected) is denoted between conditions determined by pairwise contrasts. *** = $p < .001$, ** = $p < .01$, * = $p < .05$, n.s = $p > .05$.

*Proactive Behavioural Index*

Proactive versus reactive control did not show a state-dependent shift after TMS. For PBI of RT, there was no significant main effect of block ($p$ = .840), behavioural state during TMS ($p$ = .457), session number ($p$ = .066), nor an interaction of block with behavioural state during TMS ($p$ = .612). PBI of error rates decreased as a function of block ($M_{Pre}$ = 0.77, $SD_{Pre}$ = 0.32; $M_{Post}$ = 0.61, $SD_{Post}$ = 0.54), $F_{(1, 67.73)}$ = 4.57, $p$ = .036, $R^2$ = .009. This may be driven by practice effects, as PBI of error rates also decreased across sessions ($M_{session1}$ = 0.80, $SD_{session1}$ = 0.36; $M_{session2}$ = 0.58, $SD_{session2}$ = 0.50), $F_{(1, 69.91)}$ = 7.95, $p$ = .006, $R^2$ = .070. Importantly, the engagement of proactive/reactive control was not changed as a function of behavioural state during TMS ($p$ = .607), nor as an interaction of block with behavioural state during TMS ($p$ = .477).

*Task-switching experiment*

Our within-subjects design used TOPX performance before and after TMS as a downstream readout of the function of cognitive control circuitry. With this design, it is possible that change in behavioural performance could be attributed to task switching costs or practice effects (i.e., TOPX-TOPX-TOPX in the cognitive control condition; TOPX-Perceptual-TOPX in the perceptual condition). To rule this out, we applied our experimental paradigm without administering TMS and tested for change in cognitive control performance across the first and third runs of TOPX as a function of whether a task switch occurred.

Based on the results of Experiment One, we ran a power analysis to determine the minimum required sample size to detect the smallest observed behavioural effect (D'-Context). The observed mean and standard deviation for each condition were used as generating parameters for a Monte Carlo simulation. For a grid of candidate sample sizes (N = 2 – 25 participants per condition), we generated synthetic datasets, and tested for a significant effect of task switching, repeating the process 1000 times per candidate N. Power was estimated as the proportion of simulations in which the effect of task switching was significant at $\alpha$ = .05. This indicated that a minimum of 14 participants was required to achieve 80% power. We recruited 16 participants for Experiment Two, oversampling to account for potential attrition. Participants were recruited from the online platform Prolific. Although no TMS was administered, we applied the same screening criteria as Experiment One to maintain consistency between samples.

Data collection for Experiment Two was performed in November of 2025. We highlight the consideration of the use of generative artificial intelligence (AI) among research participants in online research platforms, which has grown since the release of services such as ChatGPT in 2022 (Zhang et al., 2025). Research methods employing survey, Likert-style, or qualitative responses are particularly vulnerable to disingenuous research participation (Heyman & Heyman, 2024; Traylor, 2025; Veselovsky et al., 2025; Westwood, 2025; Zhang et al., 2025). In contrast, there is currently less evidence that generative AI can perform consistently with human response patterns in behavioural tests of cognition (Loconte et al., 2024). In particular, behavioural patterns such as conflict and stimulus discrimination effects would be difficult to emulate in real-time without direct integration into the task environment. Moreover, we employed less-conventional behavioural checks (such as testing for continued use of headphones throughout the experiment) that would be difficult to emulate without prior knowledge. Thus, we reason that it is unlikely that our online sample was contaminated by automated responding, although we emphasise the importance of continued scrutiny in online research platforms (McInnes et al., 2025).

Experiment Two followed the same procedures as the initial experiment, except that no TMS was administered. Participants each completed two sessions which involved three task runs, with a one-week washout period between sessions. The experimental tasks were administered online using PsychoPy/JS (Peirce, 2007). Parameters for the TOPX and perceptual tasks remained the same as Experiment One, but there was an enforced three-minute rest period between each task run. To maintain consistency with the previous experiment, participants were required to wear headphones, so that the masking white noise used in Experiment One could be delivered throughout the experiment. To ensure compliance, prior to each block of 30 trials, a generated audio clip instructed participants to press a randomly selected key. The experiment did not proceed without the correct input. No participants were excluded due to failing this compliance criterion. We followed the same analysis procedures as Experiment One for the behavioural data in Experiment Two. Given that we expected no effect of task switching, which relies on confirmation of the null hypothesis, to complement the frequentist tests we ran a series of Bayesian linear mixed models on the derived behavioural metrics. We report $BF_{01}$ values to evaluate the degree of evidence for the null over the alternative hypothesis and interpreted these based on the criteria outlined by Jeffreys (2006).

<u>Behavioural Change is not Explained by Task Switching Costs</u>. In Experiment Two, there was no evidence of a systematic change in manifest behaviour as a function of task switching. The interaction of TOPX block number (1, 3) with task sequence (switch, no-switch) was not significant in the frequentist linear mixed model of accuracy ($F_{(1, 45.18)} = 0.90$, $p = .349$, $R^2 = .004$), D′-Context ($F_{(1, 45.07)} = 0.03$, $p = .857$, $R^2 = .00$), or probe RTs ($F_{(1, 7653.8)} = 0.21$, $p = .650$, $R^2 = .001$). In support of this, Bayesian linear models indicated strong- to very-strong evidence (Jeffreys, 2006) against an effect of task switching on the change of TOPX accuracy ($BF_{01} = 38.00$), D′-Context ($BF_{01} = 42.33$), and probe RTs ($BF_{01} = 25.76$). However, the linear mixed model of evidence accumulation bias (*z*) indicated a significant interaction of block with task sequence, $F_{(1, 106.06)} = 16.01$, $p < .001$, $R^2 = .08$ (**Figure S2**). We further examined the DDM bias coefficient *z* in the third TOPX run as a ratio of bias in the first. Bias towards the correct response for target (A) cues was significantly enhanced (ratio > 1) when there was a task switch ($M = 1.03$, $SD = 0.05$, $p = .012$), but decreased when there was no task switch (ratio < 1, $M = 0.92$, $SD = 0.06$, $p < .001$). For non-target (B) cues, bias ratios showed no significant change when there was a task switch ($M = 1.01$, $SD = 0.05$, $p = .396$), but decreased when there was no task switch ($M = 0.85$, $SD = 0.03$, $p < .001$). Given that the direction of the effects are opposite from what we observed in our TMS experiment, it is unlikely that a confounding effect of task switching can explain the state dependent effects of TMS on decisional bias observed in Experiment One.

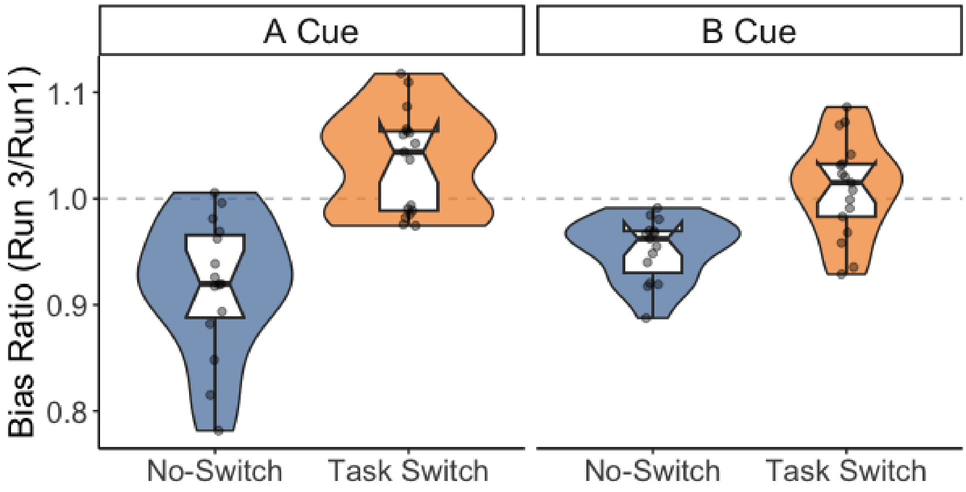


**Figure S2.** Change in evidence bias as a function of task-switching or practice effects does not explain the modification of bias by TMS observed in Experiment One.

*Stimulus-locked theta power perturbation by TMS is not explained by sustained background activity*

We considered whether the observed reductions in cue- and probe- locked theta ERSP power could be explained by changes in sustained activity, rather than an alteration of stimulus-evoked dynamics. Specifically, because computation of ERSPs involves removal of baseline activity, a sustained elevation in background power could appear as a reduction in stimulus-locked theta when the pre-stimulus baseline is removed. In other words, if ongoing activity increases, the relative change at stimulus onset may be attenuated even if evoked responses are unchanged. To assess this, we examined sustained oscillatory activity by computing power spectral densities of the baseline periods used during the computation of ERSPs. Given that spectral power reflects contributions of both rhythmic (periodic) and broadband (aperiodic) components (Donoghue et al., 2020), we considered both periodic and background aperiodic activity separately by applying spectral parameterisation via the *specparam* function in Python. Because we were primarily interested in global sustained background activity, we averaged the power spectrum across all electrodes. One participant was marked as an extreme outlier and was removed from this analysis.

Aperiodic power shifted positively after active state TMS ($p$ = .007) but not after control stimulation ($p$ = .573; **Figure S3**). In contrast, periodic power in the theta band (4 – 8 Hz) showed no change after TMS ($p$ = .131) and was not differentially modified by TMS as a function of behavioural state ($p$ = .586). Critically, if the observed reductions in ERSP theta power were driven by shifts in sustained background activity, then individuals showing larger aperiodic increases should also exhibit a larger ERSP attenuation, and the two should thus be correlated. However, we did not observe this relationship. There was no correlation between the attenuation of target cue ($r$ = .19, $p$ = .412), distractor cue ($r$ = -.39, $p$ = .169), or target probe ($r$ = .18, $p$ = .459) ERSPs and the magnitude of the shift of the accompanying aperiodic power spectra. Thus, the data do not support an argument that the observed theta modulations are explained by sustained changes in periodic or background oscillatory activity. Rather, they support the conclusion that TMS more specifically modulated stimulus-locked theta dynamics.

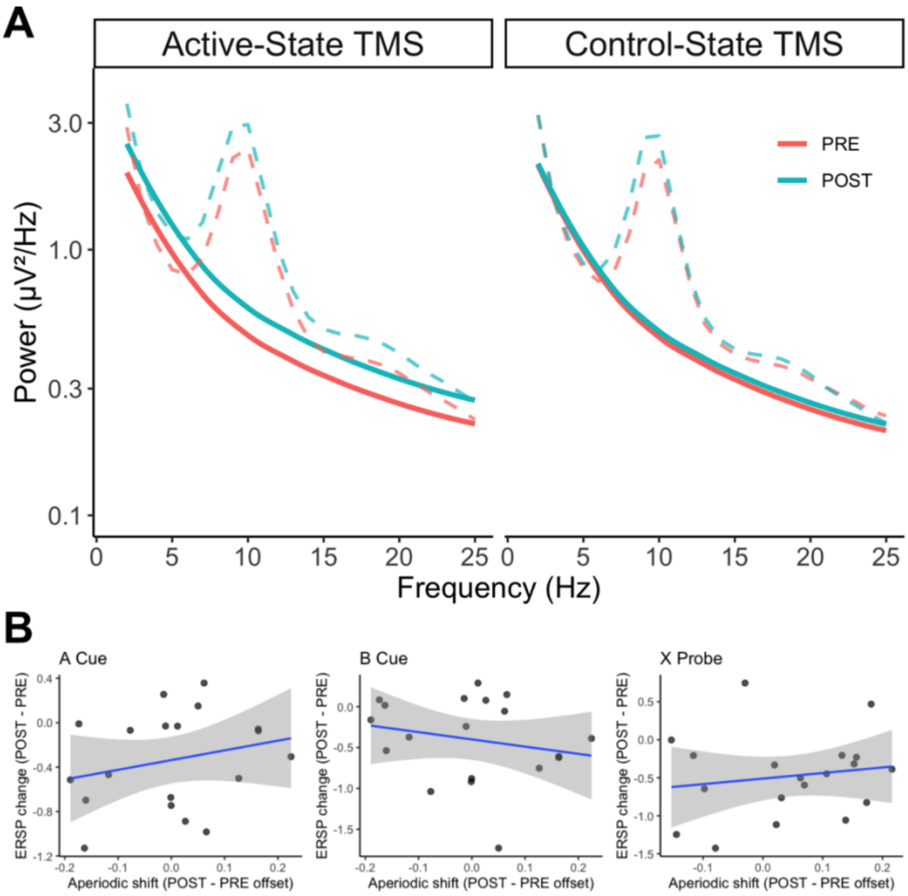


**Figure S3.** Decomposed power spectra isolated into rhythmic (periodic) and broadband (aperiodic) components, computed from the pre-stimulus baseline window used for ERSP normalisation in Experiment One. (A) Aperiodic power shifted positively after active-state TMS, but not control-state TMS. This suggests global network activation of non-oscillatory neural activity was enhanced after active-state TMS. Solid lines depict the mean aperiodic fit, dashed lines represent the average rhythmic component of the power spectra. (B) Changes in task-evoked ERSPs for target cues (A Cue), distractor cues (B Cue), and target probes (X Probe) are not explained by sustained changes in background oscillatory activity.

References

Beam, W., Borckardt, J. J., Reeves, S. T., & George, M. S. (2009). An efficient and accurate new method for locating the F3 position for prefrontal TMS applications. *Brain Stimulation*, *2*(1), 50–54. https://doi.org/10.1016/j.brs.2008.09.006

Benjamini, Y., & Hochberg, Y. (1995). Controlling the false discovery rate: A practical and powerful approach to multiple testing. *Journal of the Royal Statistical Society: Series B (Methodological)*, *57*(1), 289–300. https://doi.org/10.1111/j.2517-6161.1995.tb02031.x

Biabani, M., Fornito, A., Goldsworthy, M., Thompson, S., Graetz, L., Semmler, J. G., Opie, G. M., Bellgrove, M. A., & Rogasch, N. C. (2024). Characterising the contribution of auditory and somatosensory inputs to TMS-evoked potentials following stimulation of prefrontal, premotor, and parietal cortex. *Imaging Neuroscience*, *2*, imag–2–00349. https://doi.org/10.1162/imag_a_00349

Braver, T. S., Paxton, J. L., Locke, H. S., & Barch, D. M. (2009). Flexible neural mechanisms of cognitive control within human prefrontal cortex. *Proceedings of the National Academy of Sciences*, *106*(18), 7351–7356. https://doi.org/10.1073/PNAS.0808187106

Cohen, J. D., Perlstein, W. M., Braver, T. S., Nystrom, L. E., Noll, D. C., Jonides, J., & Smith, E. E. (1997). Temporal dynamics of brain activation during a working memory task. *Nature*, *386*(6625), 604–608. https://doi.org/10.1038/386604a0

Cohen, J. D., & Servan-Schreiber, D. (1992). Context, cortex, and dopamine: A connectionist approach to behavior and biology in schizophrenia. *Psychological Review*, *99*(1), 45–77. https://doi.org/10.1037/0033-295X.99.1.45

Donoghue, T., Haller, M., Peterson, E. J., Varma, P., Sebastian, P., Gao, R., Noto, T., Lara, A. H., Wallis, J. D., Knight, R. T., Shestyuk, A., & Voytek, B. (2020). Parameterizing neural power spectra into periodic and aperiodic components. *Nature Neuroscience*, *23*(12), 1655–1665. https://doi.org/10.1038/s41593-020-00744-x

Gelman, A., & Rubin, D. B. (1992). Inference from Iterative Simulation Using Multiple Sequences. *Statistical Science*, *7*(4), 457–472. https://doi.org/10.1214/ss/1177011136

Habibollahi Saatlou, F., Rogasch, N. C., McNair, N. A., Biabani, M., Pillen, S. D., Marshall, T. R., & Bergmann, T. O. (2018). MAGIC: An open-source MATLAB toolbox for external control of transcranial magnetic stimulation devices. *Brain Stimulation*, *11*(5), 1189–1191. https://doi.org/10.1016/j.brs.2018.05.015

Heyman, T., & Heyman, G. (2024). The impact of ChatGPT on human data collection: A case study involving typicality norming data. *Behavior Research Methods*, *56*(5), 4974–4981. https://doi.org/10.3758/s13428-023-02235-w

Jeffreys, H. (2006). Theory of probability. In *Community Care* (Issue 1638). Oxford University Press. https://doi.org/10.2307/3028701

Kleiner, M., Brainard, D., Pelli, D., Ingling, A., Murray, R., & Broussard, C. (2007). What's new in psychtoolbox-3. *Perception*, *36*(14), 1–16.

Loconte, R., Orrù, G., Tribastone, M., Pietrini, P., & Sartori, G. (2024). Challenging large language models' "intelligence" with human tools: A neuropsychological investigation in Italian language on prefrontal functioning. *Heliyon*, *10*(19), e38911. https://doi.org/10.1016/j.heliyon.2024.e38911

Ly, J. Q. M., Gaggioni, G., Chellappa, S. L., Papachilleos, S., Brzozowski, A., Borsu, C., Rosanova, M., Sarasso, S., Middleton, B., Luxen, A., Archer, S. N., Phillips, C., Dijk, D.-J., Maquet, P., Massimini, M., & Vandewalle, G. (2016). Circadian regulation of human cortical excitability. *Nature Communications*, *7*(1), 11828. https://doi.org/10.1038/ncomms11828

McInnes, A. N., Sullivan, C. R. P., Angus W. MacDonald, I. I. I., & Widge, A. S. (2025). Psychometric Validation and Preliminary Clinical Correlation of an Experiential Foraging Task. *Assessment*. https://doi.org/10.1177/10731911251376214

Peirce, J. W. (2007). PsychoPy—Psychophysics software in Python. *Journal of Neuroscience Methods*, *162*(1–2), 8–13. https://doi.org/10.1016/j.jneumeth.2006.11.017

Reimer, A. E., Dastin-van Rijn, E. M., Kim, J., Mensinger, M. E., Sachse, E. M., Wald, A., Hoskins, E., Singh, K., Alpers, A., Cooper, D., Lo, M.-C., De Oliveira, A. R., Simandl, G., Stephenson, N., & Widge, A. S. (2024). Striatal stimulation enhances cognitive control and evidence processing in rodents and humans. *Science Translational Medicine*, *16*(778), eadp1723. https://doi.org/10.1126/scitranslmed.adp1723

Rossini, P. M., Barker, A. T., Berardelli, A., Caramia, M. D., Caruso, G., Cracco, R. Q., Dimitrijević, M. R., Hallett, M., Katayama, Y., Lücking, C. H., Maertens de Noordhout, A. L., Marsden, C. D., Murray, N. M. F., Rothwell, J. C., Swash, M., & Tomberg, C. (1994). Non-invasive electrical and magnetic stimulation of the brain, spinal cord and roots: Basic principles and procedures for routine clinical application. Report of an IFCN committee. *Electroencephalography and Clinical Neurophysiology*, *91*(2), 79–92. https://doi.org/10.1016/0013-4694(94)90029-9

Satterthwaite, F. E. (1946). An Approximate Distribution of Estimates of Variance Components. *Biometrics Bulletin*, *2*(6), 110–114. https://doi.org/10.2307/3002019

Servan-Schreiber, D., Cohen, J. D., & Steingard, S. (1996). Schizophrenic Deficits in the Processing of Context: A Test of a Theoretical Model. *Archives of General Psychiatry*, *53*(12), 1105–1112. https://doi.org/10.1001/archpsyc.1996.01830120037008

Shen, C., Calvin, O. L., Rawls, E., Redish, A. D., & Sponheim, S. R. (2024). Clarifying Cognitive Control Deficits in Psychosis via Drift Diffusion Modeling and Attractor Dynamics. *Schizophrenia Bulletin*, sbae014. https://doi.org/10.1093/schbul/sbae014

Traylor, F. (2025). The threat of AI chatbot responses to crowdsourced open-ended survey questions. *Energy Research & Social Science*, *119*, 103857. https://doi.org/10.1016/j.erss.2024.103857

Veselovsky, V., Horta Ribeiro, M., Cozzolino, P. J., Gordon, A., Rothschild, D., & West, R. (2025). Prevalence and Prevention of Large Language Model Use in Crowd Work. *Commun. ACM*, *68*(3), 42–47. https://doi.org/10.1145/3685527

Westwood, S. J. (2025). The potential existential threat of large language models to online survey research. *Proceedings of the National Academy of Sciences*, *122*(47), e2518075122. https://doi.org/10.1073/pnas.2518075122

Widge, A. S., Zorowitz, S., Basu, I., Paulk, A. C., Cash, S. S., Eskandar, E. N., Deckersbach, T., Miller, E. K., & Dougherty, D. D. (2019). Deep brain stimulation of the internal capsule enhances human cognitive control and prefrontal cortex function. *Nature Communications*, *10*(1), 1536. https://doi.org/10.1038/s41467-019-09557-4

Wiecki, T. V., Sofer, I., & Frank, M. J. (2013). HDDM: Hierarchical Bayesian estimation of the Drift-Diffusion Model in Python. *Frontiers in Neuroinformatics*, *7*(JULY 2013). https://doi.org/10.3389/FNINF.2013.00014

Zhang, S., Xu, J., & Alvero, A. (2025). Generative AI Meets Open-Ended Survey Responses: Research Participant Use of AI and Homogenization. *Sociological Methods & Research*, *54*(3), 1197–1242. https://doi.org/10.1177/00491241251327130
